# Dynamic DNA methylation turnover at the exit of pluripotency epigenetically primes gene regulatory elements for hematopoietic lineage specification

**DOI:** 10.1101/2023.01.11.523441

**Authors:** Aled Parry, Christel Krueger, Tim Lohoff, Steven Wingett, Stefan Schoenfelder, Wolf Reik

**Author notes:** These authors contributed equally.

## Abstract

Epigenetic mechanisms govern developmental cell fate decisions, but how DNA methylation coordinates with chromatin structure and three-dimensional DNA folding to enact cell-type specific gene expression programmes remains poorly understood. Here, we use mouse embryonic stem and epiblast-like cells deficient for 5-methyl cytosine or its oxidative derivatives (5-hydroxy-, 5-formyl- and 5-carboxy-cytosine) to dissect the gene regulatory mechanisms that control cell lineage specification at the exit of pluripotency. Genetic ablation of either DNA methyltransferase (*Dnmt*) or Ten-eleven-translocation (*Tet*) activity yielded largely distinct sets of dysregulated genes, revealing divergent transcriptional defects upon perturbation of individual branches of the DNA cytosine methylation cycle. Unexpectedly, we found that disrupting DNA methylation or oxidation interferes with key enhancer features, including chromatin accessibility, enhancer-characteristic histone modifications, and long-range chromatin interactions with putative target genes. In addition to affecting transcription of select genes in pluripotent stem cells, we observe impaired enhancer priming, including a loss of three-dimensional interactions, at regulatory elements associated with key lineage-specifying genes that are required later in development, as we demonstrate for the key hematopoietic genes *Klf1* and *Lyl1*. Consistently, we observe impaired transcriptional activation of blood genes during embryoid body differentiation of knockout cells. Our findings identify a novel role for the dynamic turnover of DNA methylation at the exit of pluripotency to establish and maintain chromatin states that epigenetically prime enhancers for later activation during developmental cell diversification.

**Highlights:**

- We perform a detailed epigenetic characterisation of the mouse embryonic stem cell (ESC) to epiblast-like cell (EpiLC) transition in wild type, *Tet* triple-knockout (TKO) and *Dnmt* TKO lines and develop a novel clustering approach to interrogate the data.
- *Tet* TKO reduces H3K4me1 and H3K27ac levels across enhancer elements upon pluripotency exit whilst *Dnmt* TKO affects only H3K4me1 levels, suggesting a novel role for oxidative derivatives in H3K4me1 deposition.
- *Tet* TKO and *Dnmt* TKO affect enhancer priming in EpiLCs which is associated with failure to upregulate hematopoietic genes upon differentiation.
- Long-range chromosomal interactions between primed enhancers and their target genes are weakened in both *Dnmt* and *Tet* TKO.

## Introduction

The early preimplantation epiblast (∼embryonic day E3.75-E4.5 in mice) is populated by stem cells that are able to self-renew but lack the capacity to differentiate into the major cell lineages ^1,2^. In order to acquire specific cell fates during development, epiblast cells are thought to exit this “naïve” state of pluripotency and progress towards a “formative” state where the transcriptional, epigenetic, and metabolic landscape is remodelled in a way that cells acquire multi-lineage competence (in the E5.5 - E6.0 post-implantation epiblast in mice) ^3^. This transition from naïve to formative pluripotency can be modelled *in vitro* by transitioning naïve mouse embryonic stem cells (ESCs) from media containing leukaemia inhibitory factor (LIF), MAPK inhibitor and GSK3 inhibitor (known as 2i+LIF conditions) to media containing FGF2 and Activin A, which promotes the formation of epiblast-like cells (EpiLCs). EpiLCs closely resemble cells in a formative state *in vivo* ^4,5^, and they acquire germ line competence that is missing in the early epiblast and naïve ESCs ^4,6^.

DNA methylation (DNAme) is an important epigenetic mark that is dramatically remodelled during this developmental period ^7^. In mammals, DNAme principally occurs at the 5^th^ position of the base cytosine in the context of CpG dinucleotides, generating 5-methyl-cytosine (5mC). Found at high levels in the gametes, DNAme is depleted in the zygote following fertilisation but is subsequently re-established around implantation. Genome-wide levels increase from ∼25% in the early preimplantation epiblast to ∼75% in the post-implantation epiblast ^7^. Whilst these high global levels are then sustained in the majority of somatic tissues following differentiation, the precise distribution of DNAme is highly variable between tissues and cell types, especially at regulatory elements such as enhancers ^8–11^.

The molecular machinery that deposits and removes DNAme are relatively well understood: DNA methyltransferases 3A and 3B (DNMT3A/B) deposit DNAme *de novo* and DNMT1 maintains DNAme patterns following cell division (together with cofactors) ^12^. Conversely, DNAme can be lost passively when the levels or activity of the DNMTs are reduced or can alternatively be removed in an active process involving sequential oxidation by Ten Eleven Translocation enzymes (TET1, 2, 3) to 5-hydroxy-, 5-formyl- and 5-carboxy-cytosine (5hmC, 5fC and 5caC respectively) ^13–16^. These derivatives are not efficiently recognised by DNMT1 and are therefore lost upon replication ^17–19^. Alternatively, 5fC and 5caC can also be removed in an active and cell-cycle independent manner by thymine-DNA glycosylase and base excision repair ^15,20^.

As cells exit naive pluripotency, both DNMTs and TETs are paradoxically co-expressed at high levels and target overlapping genomic regions leading to a cyclical turnover of methylation. This DNAme turnover occurs genome-wide but is especially prevalent across regulatory elements such as enhancers ^21–24^. Whilst the role of DNAme turnover is not clear, the remodelling of DNAme in early development is functionally critical. Embryonic stem cells lacking all three DNMTs or TETs (triple knockout cells, TKO) remain pluripotent but they do not differentiate effectively ^25,26^. *Dnmt1* or Dnmt*3b* knockout embryos and *Tet* TKO embryos die shortly after gastrulation with severe but poorly characterised defects including abnormal primitive streak patterning and differentiation of mesodermal tissues ^27–29^. Mutations in both *Dnmt* and *Tet* genes have been observed frequently in haematological malignancies and clonal hematopoiesis ^30–36^, and more recently lineage specific knockouts and chimeric mice have demonstrated that TET enzymes are required for efficient hematopoiesis ^37,38^, suggesting an important role for methylation and oxidation in this lineage. Bulk RNA-sequencing and whole genome bisulfite-sequencing (WGBS) of *Tet* TKO embryos implicated *Lefty1/2* hypermethylation and perturbed NODAL and WNT signalling as a driver of these defects ^28,39,40^, but the broader effect of TET activity loss on the histone modification landscape and enhancer activity was not investigated, despite enhancers being primary regions of DNAme turnover.

In contrast to active enhancers (marked by H3K4me1 and H3K27ac) that drive expression of associated genes, primed or poised enhancers (marked by H3K4me1 alone or a combination of H3K4me1 and H3K27me3, respectively) are silent but molecularly prepared for gene activation upon stimulation (e.g. following differentiation) ^41–43^. Recent multi-omic data has demonstrated that enhancers driving ectoderm specific gene expression programmes are already hypomethylated and accessible in the epiblast, suggesting priming, whilst mesoderm and endoderm enhancers are not primed at this stage ^10^. However, the role of DNA methylation and oxidation in regulating this epigenetic environment is unknown.

Here, we profile in depth the transcriptome, chromatin structure and epigenome of mouse ESCs across culture conditions that model the exit of pluripotency, and we develop a novel analysis approach that uses promoter-capture HiC data (PCHi-C) to identify promoter-regulatory element pairs. To understand the effects of DNAme turnover on this transcriptional and epigenetic landscape, we perform the same molecular characterisation in cells lacking DNAme (*Dnmt* TKO) or oxidation (*Tet* TKO). Excitingly, we find that both knockouts result in reduced H3K4me1 levels whilst H3K27ac is only affected upon *Tet* TKO suggesting a novel role for methylation turnover in H3K4me1 deposition. Gene expression changes upon *Tet* TKO were linked to enhancer number, where highly expressed genes interacting with multiple enhancers were more likely to be downregulated than those with fewer putative enhancer interactions. Loss of methylation and oxidation result in distinct transcriptional defects upon differentiation into embryoid bodies (EBs): *Dnmt* TKO cells fail to exit pluripotency effectively whilst *Tet* TKO EBs have distinct lineage biases including defects in blood specification. Finally, we describe extensive loss of enhancer priming at regulatory elements associated with key hematopoietic transcription factors in both knockouts at the EpiLC stage, which may explain the failure to upregulate this transcriptional programme upon differentiation.

## Results

### EpiLC establishment is accompanied by epigenetic remodelling

To understand the interplay between higher-order chromatin structure and epigenetic chromatin modifications, and how this governs the transcriptional programmes that underpin formative pluripotency, we transitioned wild type (WT) naïve ESCs (cultured in 2i-LIF conditions) to EpiLCs ^4^ and profiled chromatin states using a broad range of genomics approaches. These included RNA-sequencing, chromatin immunoprecipitation and sequencing (ChIP-seq), assay for transposase accessible chromatin and sequencing (ATAC-seq), whole genome bisulfite sequencing (WGBS), 5hmC pulldown and sequencing (HMCP-seq), and PCHi-C **(Fig. 1a)**.

**Figure 1.**
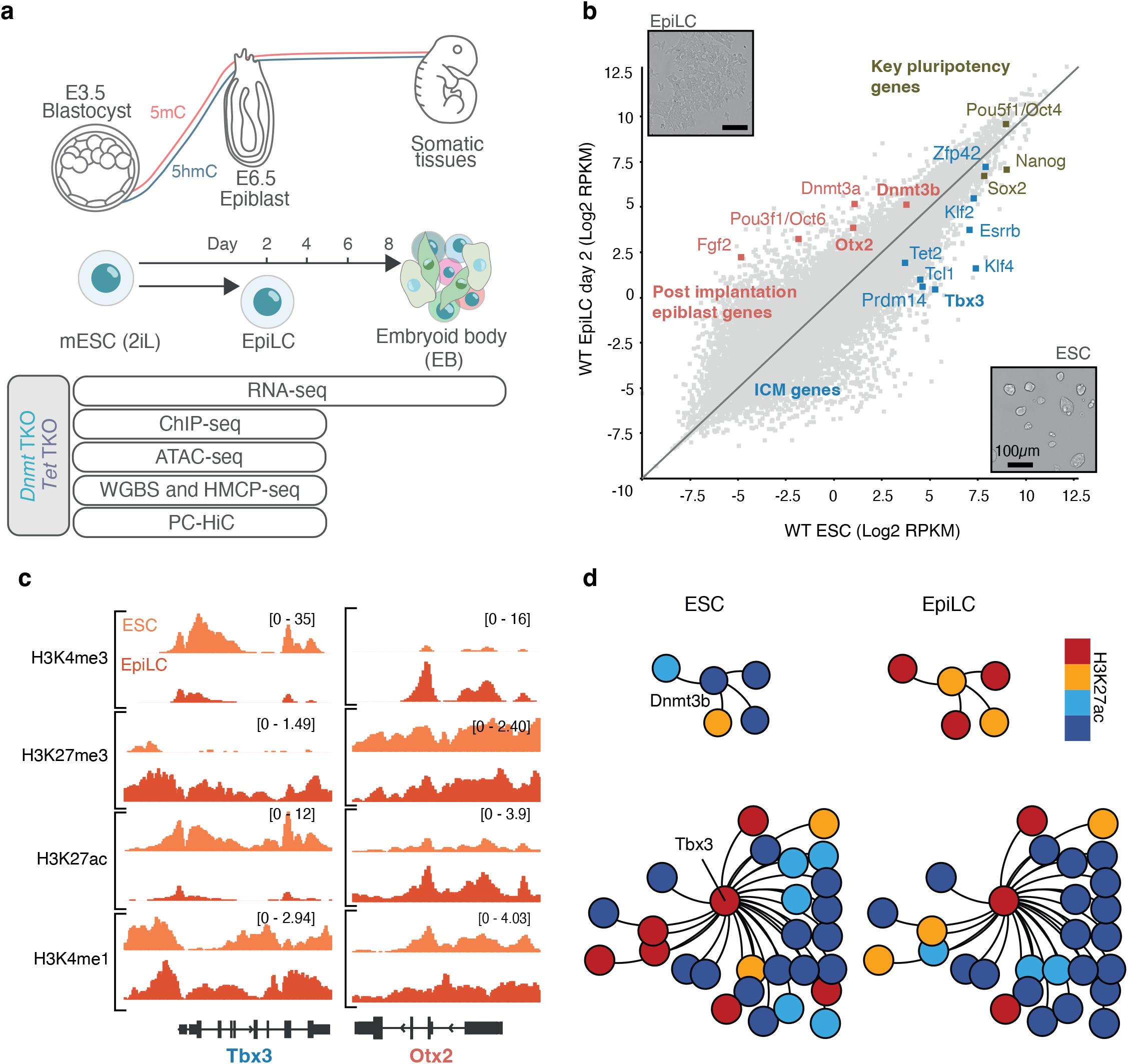
EpiLC establishment is accompanied by epigenetic remodelling **(a)** Diagram illustrating the mouse embryonic stem cell (ESC) to epiblast-like cell (EpiLC) transition and an EB differentiation time course performed in this study, together with the *in vivo* stages that these represent (top). Datasets collected and genotypes interrogated in this study are indicated (bottom). **(b)** Scatter plot showing relative levels of mRNA (Log_2_ reads per kilobase of sequence per million reads - RPKM) in wild type (WT) ESCs against WT EpiLCs. Select genes associated with the inner cell mass (ICM, blue), post-implantation epiblast (red) and pluripotency (brown) are labelled. Representative images of the cells are presented as insets (scale bar = 100*µm*). **(c)** Representative genome browser images showing ChIP-seq data at the promoters of a gene downregulated (*Tbx3*) and upregulated (*Otx2*) upon ESC to EpiLC transition. RefSeq gene annotations are included **(d)** Force directed graph of promoters and promoter interacting regions (PIRs) (nodes) connected by the interactions between them (edges) generated using Promoter Capture Hi-C (PCHi-C) data. Subclusters centred on genes of interest (*Dnmt3b* and *Tbx3*) and their PIRs coloured by the levels of H3K27ac in ESCs and EpiLCs (red = high, blue = low, for thresholding criteria see accompanying material) **(b-d)** WT cells represented are matched to *Tet* TKO cells used in following experiments. Values are the average of 3 biological replicates.

Over a period of 48 hours cells became morphologically flatter and began expressing key post-implantation associated genes including *Pou3f1, Fgf2, Otx2* and *Dnmt3a/b* **(Fig. 1b)**. Naïve pluripotency genes including *Tbx3, Klf4* and *Esrrb* were also robustly downregulated whilst core pluripotency genes such as *Oct4* and *Sox2* remained expressed at high levels **(Fig. 1b)**. More broadly, we identified 434 significantly upregulated genes and 765 significantly downregulated genes (DESeq p <0.05 and dynamic fold-change filter) that were enriched in gene ontology (GO) terms such as MAPK signalling and focal adhesion, respectively **(Supplemental Fig. 1a)**.

The promoters of both activated and repressed genes remained largely hypo-methylated **(Supplemental Fig. 1b)** despite the expected global accumulation of DNA methylation observed **(Supplemental Fig. 1c)**, suggesting that promoter methylation plays a minor role in gene regulation during the formation of EpiLC cells. Histone modification changes at promoters reflected expression change for some but not all genes. For example, 316 of 434 upregulated genes (∼73%) accumulated H3K4me3 (DESeq p<0.05), a mark associated with active promoter elements. Similarly, 300 of 765 downregulated genes lost H3K4me3 (∼39%) from their promoters **(Fig. 1c, Supplemental Fig. 1d)**. Similar correlations were observed for H3K27ac levels and accessibility levels **(Supplemental Fig. 1d)**.

Regulatory elements, such as enhancers, are thought to interact with cognate promoters via three-dimensional loops ^44^. To connect enhancers with their putative target genes, we used PCHi-C data to build a network ^45^ in which each node represented a genomic location (either a promoter or a promoter interacting region [PIR]) and each edge represented a PCHi-C interaction. For visualisation we used a force-directed graph layout in which highly interacting regions are pulled close together, and epigenetic information can be superimposed (similar to Canvas in ^46^, **Supplemental Fig. 2a, b**). Colouring edges by the levels of histone modifications at promoters or PIRs did not reveal substantial changes between ESC and EpiLC cells on a global scale; however, we detected clusters of active or inactive genes and regulatory elements **(Supplemental Fig. 2b)**. For example, there were distinct clusters of H3K27me3 marked genes and regulatory elements, including clusters of homeobox (*Hox*) genes ^47^ known to be repressed by Polycomb repressive complex 2 (PRC2) and PRC1 in embryonic stem cells **(Supplemental Fig. 2b)** ^48,49^.

**As anticipated, genes activated upon ESC to EpiLC transition tended to interact with a greater number of PIRs that gain the active histone modifications H3K27ac and H3K4me1, whilst repressed genes tended to interact with more PIRs that lose these marks (Supplemental Fig. 2c-f)**. For example, of the PIRs interacting with upregulated genes, 11.2% accumulated H3K27ac (change in RPKM >1 between conditions), whilst just 0.1% of PIRs interacting with downregulated genes gained H3K27ac upon ESC to EpiLC transition **(Supplemental Fig. 2f)**. A subnetwork centred on the well-characterised post implantation gene *Dnmt3b* showed that all significant PIRs detected (4/4) gained H3K27ac upon the EpiLC transition, concomitant with activation of the gene **(Fig. 1b, d)**. These PIRs are putative enhancer elements that may play a role in gene activation. Conversely, the naïve pluripotency associated gene *Tbx3* formed significant interactions with 27 different PIRs. Of these, 8 were marked by H3K27ac in ESCs, and 5 of these lost H3K27ac upon transition to the EpiLC state, consistent with transcriptional repression **(Fig. 1b, d)**. Similar networks built around other genes involved in naïve pluripotency and the post implantation epiblast showed similar patterns, where histone modifications on some but not all PIRs correlated with expression change **(Supplemental Fig. 3a, b)**. As some of these PIRs are likely gene regulatory elements such as enhancers, we next sought a method to define these systematically.

### A novel clustering approach to dissect gene regulatory element - chromatin interactions

To define promoter-PIR interactions in a systematic manner, we developed a novel method whereby we treated each promoter-PIR pair individually and used the epigenetic and transcriptional data that we had collected for each pair to perform dimensionality reduction and expectation-maximisation based clustering **(methods, Fig. 2a)**. The resulting UMAP separated promoter-PIR pairs into 20 clusters based on transcription of the associated gene and epigenetic information from both interacting regions including histone modifications, accessibility, DNAme and interaction strength **(Fig. 2b, c, Supplemental Fig. 4a)**. For example, cluster 16 consisted of active promoters (transcribed, marked by H3K4me3 and H3K27ac) interacting with active enhancers (marked by high levels of H3K27ac and H3K4me1) **(Fig. 2b-d)**. Cluster 17 consisted of inactive promoters interacting with putative “poised” enhancers (marked by HK4me1 and HK27me3) **(Fig. 2b, c)**. Clusters 6, 12 and 19 consisted of lowly or unexpressed genes interacting with H3K4me1 marked PIRs, which are putative “primed” enhancers ^41–43^. Clusters 1, 2 and 15 were marked by varying levels of H3K27me3 at the promoter, including repressed and bivalent genes, interacting with unmarked PIRs. Finally, there were a number of clusters that contained active promoters interacting with unmarked PIRs (3, 10, 13, 14, 18), as well as some clusters containing inactive promoters interacting with unmarked PIRs (0, 5, 7, 8 11).

**Figure 2.**
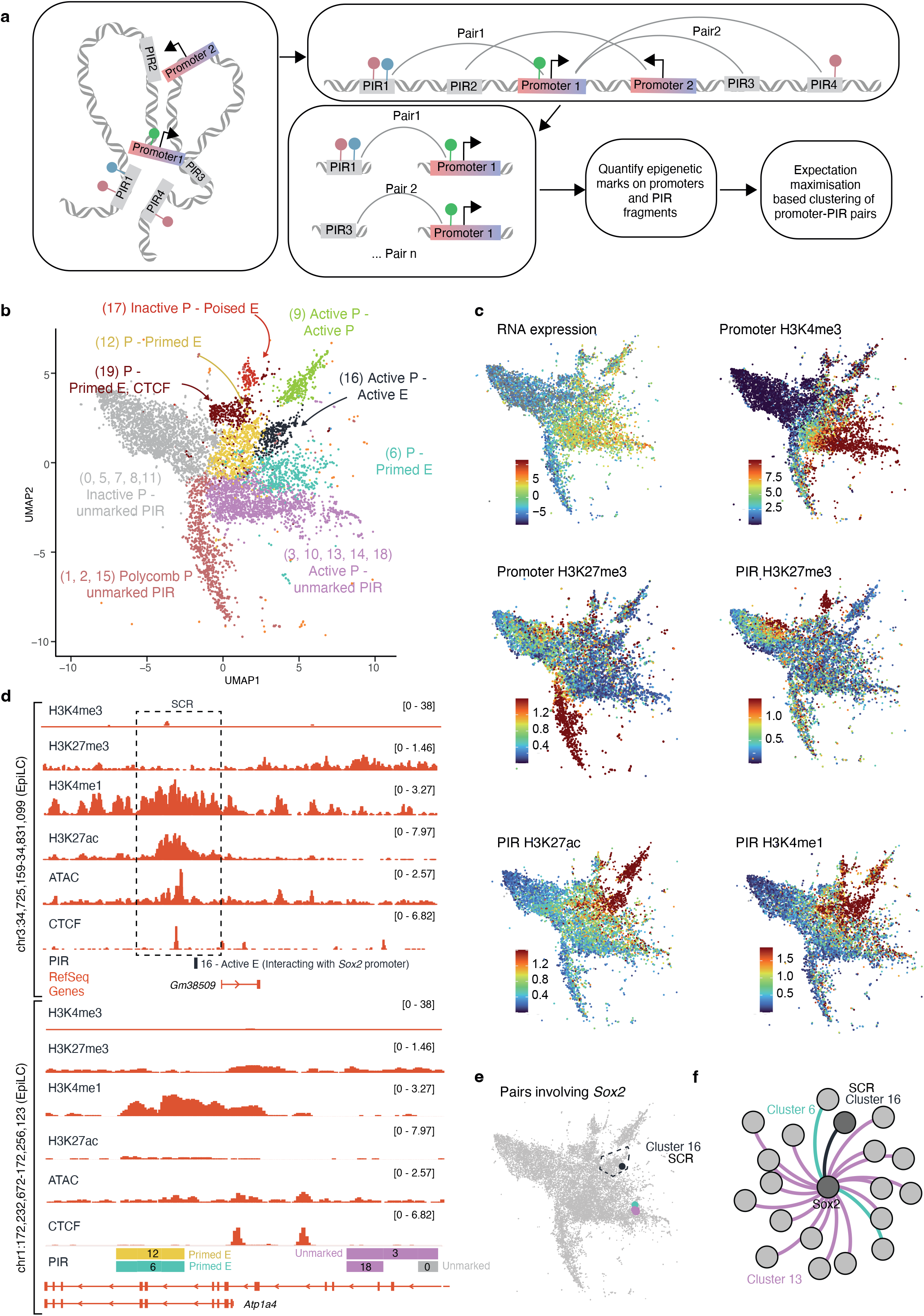
A novel clustering approach dissects the regulatory complexity of EpiLCs **(a)** Diagram illustrating the computational approach undertaken. Epigenetic, transcriptional and 3D interaction data from promoters and PIRs was quantified such that each region could be involved in multiple promoter-PIR pairs. These pairs were used for dimensionality reduction and expectation maximisation based clustering. Arrows indicate gene transcriptional start sites and lollipops represent epigenetic modifications, such as histone modifications. **(b)** Results from the computational approach described in (a) plotted using uniform manifold approximation and projection (UMAP), labelled with cluster numbers. Colours represent groupings of clusters which share similar epigenetic information on the promoter and PIR. P = Promoter, E = Enhancer. **(c)** UMAPs as in (b) coloured by the levels of transcription (at the promoter linked gene) or histone modifications at the promoters or PIRs. Colours are reads per kilobase of fragment per million (RPKM), capped at 3 median absolute deviation (MAD). **(d)** Representative genome browser images of ChIP and ATAC-seq data together with annotations generated using the clustering approach described in (a). An active enhancer, (the *Sox2* control region (SRC), top) and primed enhancer (bottom) are displayed. **(e)** UMAP as in (b) highlighting pairs involving the *Sox2* promoter. **(f)** Subnetwork showing the *Sox2* promoter and directly associated PIRs based on PCHi-C data. The interactions between the promoter and the PIRs (edges) are coloured based on the clusters defined in (b). The PIR overlapping the SCR is indicated. **(b-f)** Data represented are WT cells matched to *Tet* TKO cells used in following experiments. Values are the average of 3 biological replicates.

Interestingly, the cluster of active promoters interacting with active enhancers (cluster 16) contained the promoters for pluripotency associated genes (e.g. *Sox2, Sall1, Nanog*) that are highly expressed at the epiblast stage. Clusters of active promoters interacting with unmarked PIRs (e.g., cluster 10) frequently involved housekeeping genes, such as genes involved in mRNA processing. Polycomb marked clusters (e.g., cluster 2) were enriched for lineage-specific genes that are commonly upregulated upon differentiation (e.g. *Olig2, Hand1, Hoxb1, Neurog1)*. Meanwhile, clusters of inactive promoters interacting with unmarked PIRs (e.g., Cluster 8) contained highly lineage specific genes (such as olfactory receptors). **(Supplemental Fig. 4a)**. These results suggest fundamentally different modes of regulation for genes required at distinct times during development.

As anticipated, single genes were involved in more than one promoter-PIR pair, appearing in more than one cluster. For example, *Sox2* was involved in promoter-PIR pairs found in clusters 16 (active promoter -active enhancer), 6 (promoter - primed enhancer), and 13 (active promoter - unmarked PIR) **(Fig. 2d-f)**. Indeed, the active enhancer region identified using our method overlapped the previously described Sox2 control region (SCR), essential for *Sox2* activity ^50^.

Putative active enhancers defined using our method overlapped well with elements defined by others using ChIP-seq **(Supplemental Fig. 4b)** ^51^. Moreover, consistent with previous work, we found that polycomb marked promoters (clusters 1, 2, 15) and poised enhancers (cluster 17) dramatically accumulate H3K27me3 and form more significant Hi-C interactions upon ESC to EpiLC transition and global methylation of the genome **(Supplemental Fig. 4c-e)** ^52^. Interestingly, H3K4me1 also accumulated upon transition from ESC to EpiLC at putative active (cluster 16) and primed enhancers (clusters 6, 12 and 19) whilst H3K27ac levels were unchanged **(Supplemental Fig. 4f-h)**. Thus, our data show that some chromatin modifications are stable whilst others are re-organised during pluripotency exit.

Overall, our dataset and clustering method form a comprehensive picture of the transcriptional and epigenetic landscape of EpiLCs that (1) integrates numerous layers of molecular information, (2) clearly defines pairs of promoters and putative regulatory elements (allowing one to identify elements of particular interest such as the SCR) and (3) can be easily interrogated by, for example, filtering for different types of elements or filtering for promoter-PIR pairs involving genes of interest.

### H3K4me1 and H3K27ac are differentially affected by DNA methylation

DNA methylation (5mC) and oxidative derivatives (5hmC, 5fC, 5caC) accumulate during the transition from naïve ESC to EpiLC when *Dnmt* and *Tet* genes are co-expressed at high levels ^22^. We therefore asked whether DNA methylation and oxidation are important to establish the histone post-translational modification landscape of EpiLC cells that we have defined.

We generated epigenetic datasets (as in **Fig. 1a**) from *Dnmt* triple knockout (TKO) ^53^ and *Tet* TKO ^28^ ESCs and EpiLCs. RNA-seq analysis demonstrated that some *Tet1/2/3* and *Dnmt1/3a/3b* transcript were still present despite excision of the catalytic domains (in the *Tet* TKO line) or frame-shift mutations (in the *Dnmt* TKO line) introduced when the cells were generated **(Supplemental Fig. 5a)**. Importantly however, DNA mass spectrometry and WGBS confirmed that 5hmC was absent from both lines, whilst 5mC was absent from the *Dnmt* TKO line but accumulated in the *Tet* TKO line **(Supplemental Fig. 5b, c)**.

We first plotted the levels of H3K4me1 and H3K27ac across the active enhancer elements defined in this study (cluster 16) in TKO EpiLCs relative to matched WT cells **(Fig. 3a-d)**. Excitingly, we found that H3K4me1 was depleted in both TKO lines relative to their WT controls, whilst H3K27ac was depleted only in the *Tet* TKO line **(Fig. 3a-d)**. The same was true across primed and poised enhancers (e.g., clusters 12 and 17 respectively), though H3K27ac levels were expectedly low at these **(Fig. 3c, d)**.

**Figure 3.**
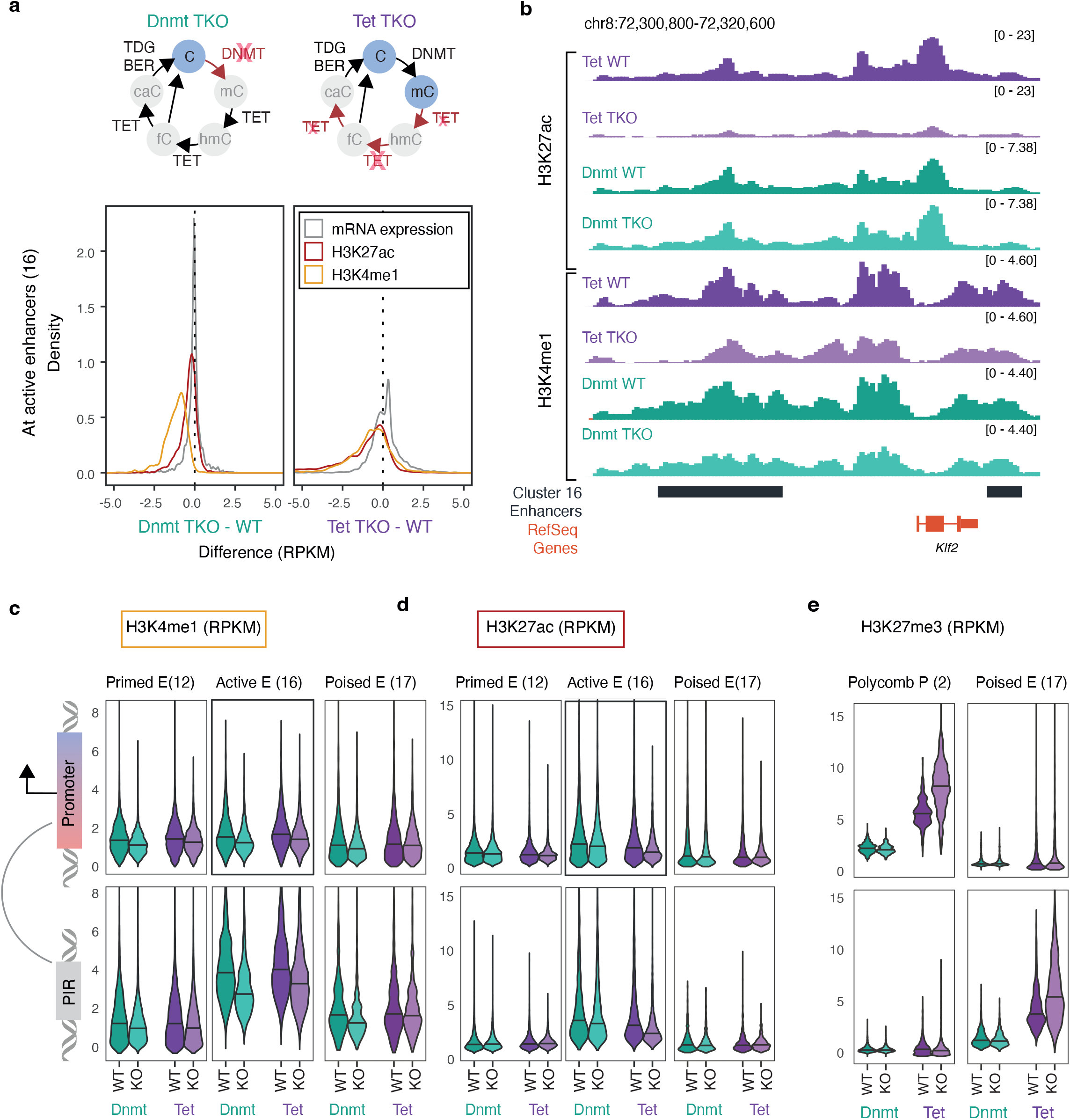
The levels of histone modifications at regulatory elements are differentially affected by DNA methylation and oxidation **(a)** Density plots showing the levels of H3K4me1 (orange), H3K27ac (red) and expression (grey) in *Dnmt* TKO EpiLCs and *Tet* TKO EpiLCs across active enhancer elements defined in Figure 2 (cluster 16 PIRs) relative to their respective WT controls. For reference the DNA modifications present or absent in each knockout line are indicated in blue and grey circles respectively (top). **(b)** Genome browser images displaying H3K4me1 and H3K27ac enrichment across active enhancers defined in Figure 2 in *Tet* and *Dnmt* TKO lines and their respective controls. **(c, d)** Violin plots displaying levels of H3K4me1 (c) and H3K27ac (d) across promoters and PIRs in clusters 12, 16 and 17 - putative primed, active and poised enhancers respectively (RPKM). **(e)** Violin plots displaying levels of H3K27me3 across promoters and PIRs in clusters 2 and 17 - polycomb marked promoters and poised enhancers, respectively (RPKM). **(a-d, f)** Values are the average of three biological replicates.

Considering the accumulation of H3K4me1 we observed during the ESC to EpiLC transition **(Supplemental Fig. 4f)** required both DNMT and TET activities **(Fig. 3a)**, our results suggest that oxidation derivatives (5hmC, 5fC or 5caC) are important for H3K4me1 deposition. Meanwhile, because H3K27ac levels are stable during the ESC to EpiLC transition **(Supplemental Fig. 4g)** and unaffected by loss of DNMT activity **(Fig. 3a)**, we reason that deposition of this mark requires (maintenance of) unmethylated DNA.

Next, given that we observed an accumulation of H3K27me3 across bivalent and repressed promoters (clusters 1, 2, 15) and poised enhancers (cluster 17) upon ESC to EpiLC transition **(Supplemental Fig. 4c, d)**, and that TET proteins are known to localise with PRC2 at bivalent promoters ^54–57^, we asked what effect loss of DNMT or TET activity has on this mark. We observed a pronounced accumulation of H3K27me3 across repressed promoters and poised enhancers in *Tet* TKO EpiLCs relative to WT controls **(Fig. 3e, Supplemental Fig. 5d)** together with a loss of H3K27me3 largely from non-promoter regions that do not feature prominently in promoter - PIR clusters **(Supplemental Fig. 5d)**. Given that these cells express Tet catalytic mutants **(Supplemental Fig. 5a)**, our results suggest that TETs facilitate H3K27me3 deposition in a manner independent of their catalytic activity, and indeed that this can be boosted in the absence of the catalytic domain (e.g., by altering PRC2 targeting or activity). This is consistent with recent publications showing that full length *Tet1* knockout cells, but not catalytic mutants, lose H3K27me3 from promoters ^58,59^.

Finally, we investigated what happens to CTCF binding upon loss of DNAme or its oxidation. We observed reduced levels of CTCF across the promoters and PIRs where it is enriched (cluster 6 promoters and cluster 19 PIRs) in *Tet* TKO cells (where DNAme is increased) but not in *Dnmt* TKO cells (where DNAme is absent) (**Supplemental Fig. 5e)**, consistent with the previously described role of DNA methylation in preventing CTCF binding ^60,61^.

### Highly expressed genes that interact with multiple enhancers are sensitive to *Tet* TKO

Enhancers are thought to drive expression of linked genes, and we therefore hypothesised that the dramatic changes at enhancer chromatin that we had observed would correlate with transcriptional downregulation of linked genes. To investigate this, we profiled mRNA expression levels in WT, *Dnmt* and *Tet* TKO EpiLCs using RNA-seq.

*Dnmt* TKO resulted in more upregulated (735) than downregulated (373) genes in EpiLC cells **(Fig. 4a)**, consistent with the role of DNA methylation in gene repression. Indeed, the majority of the upregulated genes were ESC expressed genes that failed to be repressed upon EpiLC transition (445/735 genes, **Fig. 4b)**. *Dnmt* TKO also resulted in downregulation of genes associated with MAPK signalling and pluripotency; the latter included the *Dnmt3a/b* and *Dnmt1* genes themselves. *Tet* TKO resulted in upregulation of 834 and downregulation of 576 genes in EpiLC cells, the latter of which were especially enriched in pluripotency genes by gene ontology analysis (adjusted p-value = 5.214e-10) **(Fig. 4c, d)**. The downregulated pluripotency genes included *Lefty1*, which was previously shown to be aberrantly repressed by DNAme in *Tet* TKO mice ^28^. Most of these genes were expressed at near normal levels in TKO ESCs **(Fig. 4c)**, suggesting that their downregulation is caused by the inappropriate accumulation of DNAme and/ or loss of oxidative derivatives (5hmC, 5fC, 5caC) at regulatory elements upon transition to EpiLC.

**Figure 4.**
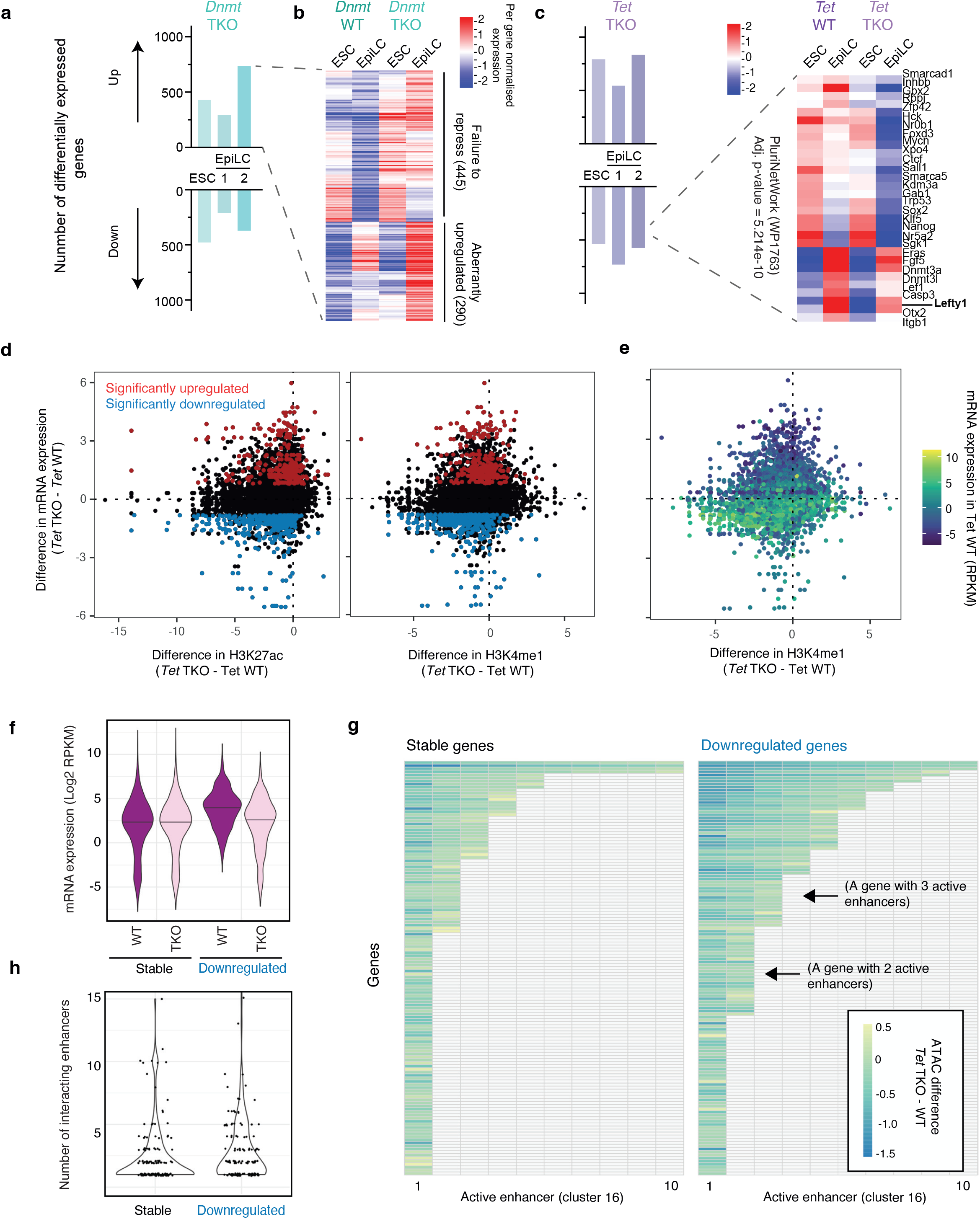
Highly expressed genes interacting with multiple enhancers are sensitive to *Tet* TKO **(a)** Number of differentially expressed genes in *Dnmt* TKO ESCs and EpiLCs following 1 or 2 days of transition. Numbers are based on DESeq and dynamic fold change filters applied to three biological replicates. **(b)** Heatmap showing per gene normalised expression of genes aberrantly upregulated in *Dnmt* TKO EpiLCs relative to matched WT controls. **(c)** Number of differentially expressed genes in *Tet* TKO ESCs and EpiLCs following 1 or 2 days of transition (left) and heatmap showing per probe normalised heatmaps of aberrantly downregulated genes from the GO term “PluriNetWork” (right). Numbers are based on DESeq and dynamic fold change filters applied to three biological replicates. **(d)** Change in H3K27ac and H3K4me1 levels at PIRs against change in mRNA expression (of the gene associated with the linked promoter fragment) in *Tet* TKO cells relative to matched WT controls across active enhancers (defined in Fig. 2). Enhancers linked to differentially expressed genes are indicated. **(e)** As in (d) except that points are coloured based on expression of enhancer linked genes in *Tet* WT cells. **(f)** mRNA expression of genes significantly downregulated in *Tet* TKO EpiLCs and a representative set of stable genes that are not differentially expressed. Downregulated genes are expressed at a higher level than stable genes in WT cells. **(g)** Heatmap showing the change in accessibility (ATAC-seq signal) at enhancer elements linked to genes described in (f). Rows represent genes and columns represent enhancers. Genes are sorted by the number of interacting enhancers (top to bottom) and enhancers are sorted by largest change (left to right). **(h)** Number of enhancers interacting with stable genes and genes downregulated in *Tet* TKO EpiLCs described in (f).

Surprisingly, we did not observe a strong negative correlation between mRNA expression change and H3K4me1/H3K27ac change at linked enhancers (cluster 16) in *Tet* TKO **(Fig. 4d, e)** or *Dnmt* TKO **(Supplemental Fig. 6a)** EpiLCs. Significantly downregulated genes had a tendency to interact with enhancers that lose active histone modifications in *Tet* TKO cells but not in *Dnmt* TKO cells. Therefore, downregulation of genes in *Dnmt* TKO EpiLCs relative to WT controls may be a result of cells failing to exit pluripotency effectively rather than failing to activate enhancers (consistent with our finding that many ESC genes are not repressed effectively - **Fig. 4a**).

We hypothesised that the lack of correlation between enhancer modification change and expression change was a result of enhancer redundancy (multiple regulatory elements controlling single genes). To test this, we studied the change in ATAC-seq, H3K4me1 and H3K27ac signal upon *Tet* TKO at the putative enhancer elements associated with significantly downregulated genes versus a representative set of stable genes that were not differentially expressed **(Fig. 4f, g, Supplemental Fig. 6b-d)**. We observed that enhancers associated with downregulated genes had a tendency to lose more ATAC-seq signal (accessibility) in the *Tet* TKO EpiLC cells than enhancers associated with stable genes **(Fig. 4g)**, despite starting with similar levels of accessibility **(Supplemental Fig. 6b)**. The same was true of H3K4me1 and H3K27ac levels across these two sets of enhancer elements **(Supplemental Fig. 6c, d)**. Interestingly, genes that were downregulated upon *Tet* TKO were more highly expressed in WT EpiLCs than genes that were stable **(Fig. 4f)** and they, on average, interacted with more enhancer elements **(Fig. 4h)**. These data suggest that highly expressed genes, which are dependent on multiple enhancer elements to maintain such expression levels, are sensitive to impaired enhancer function upon *Tet* TKO. Meanwhile, stable genes interacting with fewer enhancer elements are less affected by enhancer disruption upon *Tet* TKO.

### *Tet* TKO results in impaired upregulation of hematopoietic genes

Interestingly, many of the regulatory elements that lost active histone modifications in *Tet* TKO cells were linked to lowly expressed genes in *Tet* WT cells **(Fig. 4e)**. One possibility is that these regulatory elements prime associated genes for expression later on during development, and that loss of active histone modifications at these elements may then affect their ability to be upregulated.

To test this hypothesis, we generated RNA-seq data across an 8 day embryoid body (EB) differentiation time course (sampling every 2 days) in *Dnmt* TKO and *Tet* TKO mESCs as well as matched WT controls **(Fig. 1a)**. Principal component analysis (PCA) of the EB, ESC and EpiLC RNA-seq data separated the WT samples along PC1 indicating a gradual transcriptional change upon differentiation **(Fig. 5a, b)**. According to GO analysis, genes associated with differentiated cell types were upregulated (e.g., terms related to ossification and heart development) whilst pluripotency genes were downregulated in both WT lines **(Supplemental Fig. 7a, b)**. Many differentially expressed genes were common between the WT lines, indicating that our protocol was robust **(Supplemental Fig. 7c, d)**.

**Figure 5.**
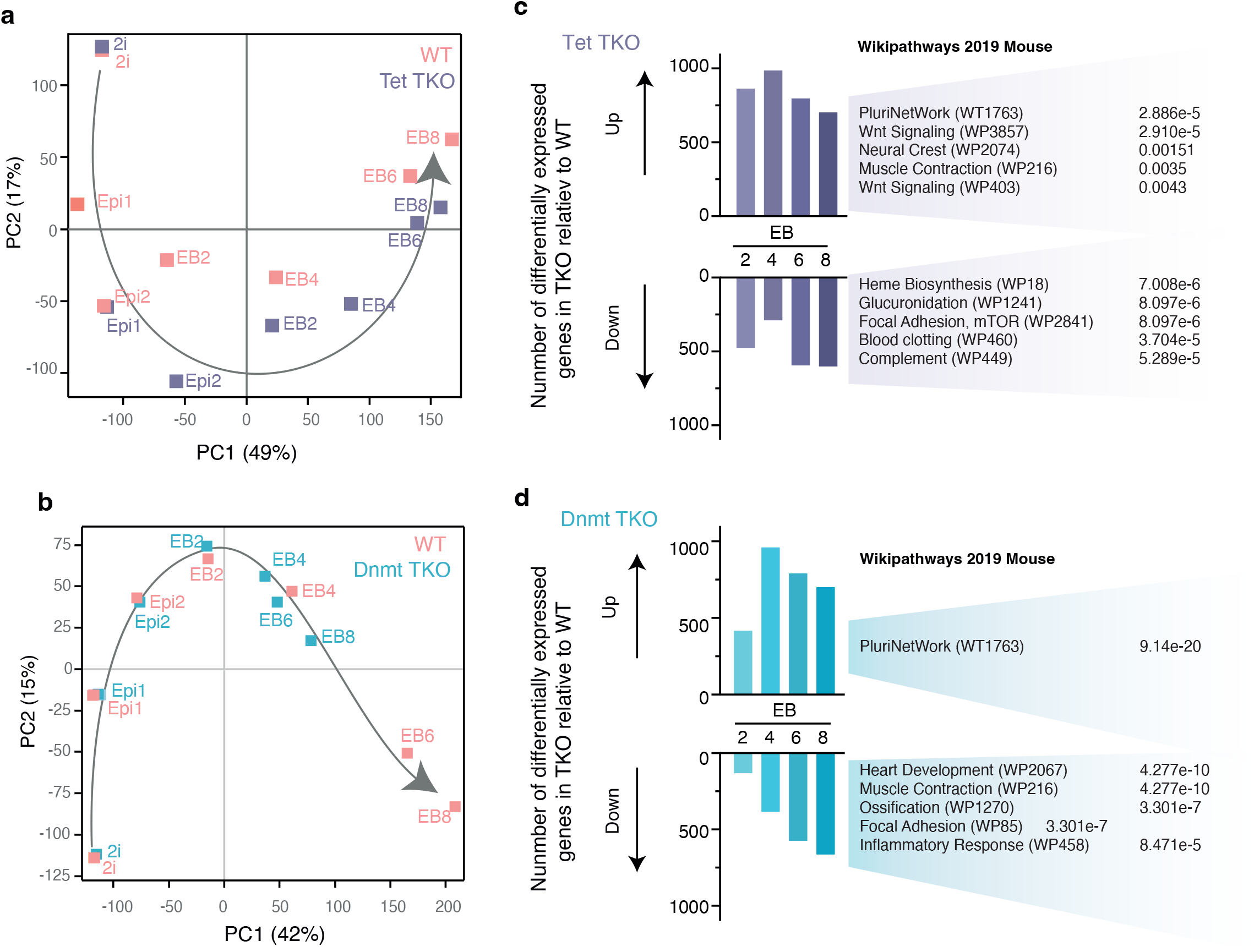
*Tet* and *Dnmt* TKO result in distinct differentiation defects **(a, b)** PCA plots of mRNA-seq data generated from WT, *Tet* TKO **(a)** and *Dnmt* TKO **(b)** ESCs (2i-LIF) transitioned to EpiLCs (Epi) or embryoid bodies (EB) for 8 days, sampling every two days (where labels indicate number of days of EpiLC transition or EB differentiation). **(c, d)** The number of differentially expressed genes that are upregulated or downregulated in *Tet* **(c)** and *Dnmt* **(d)** TKO cells relative to their respective WT controls in EBs differentiated for 2, 4, 6 or 8 days (left) and the top gene ontology categories (Wikipathways 2019 Mouse) that differentially expressed genes in day 8 EBs were found to enrich (right) including adjusted p-values.

Based on PCA, *Tet* TKO cells appeared to exit pluripotency more quickly than their WT counterparts but did not appear to reach the same endpoint **(Fig. 5a)**, suggesting transcriptional defects in both pluripotency and differentiation programmes. Meanwhile, *Dnmt* TKO had a more severe phenotype where day 8 EBs transcriptionally resembled WT EBs at day 4, suggesting a differentiation block **(Fig. 5b)**. A few hundred genes were differentially expressed in both knockout lines at the EpiLC stage and in EBs **(Fig. 5c, d)** but the limited overlap between these gene lists suggests fundamentally different transcriptional defects upon loss of DNMT and TET activity **(Supplemental Fig. 7e, f)**. Based on GO analysis, *Dnmt* TKO EBs at day 8 expressed pluripotency genes at higher levels than their WT counterparts but failed to efficiently upregulate genes associated with heart, muscle, bone and blood development, consistent with a transcriptional block **(Fig. 5d)**. Interestingly, *Tet* TKO EBs at day 8 showed lineage biases, including an upregulation of genes associated with Wnt signalling, muscle and neural crest formation, and a downregulation of genes strongly enriched in GO terms associated with the hematopoietic system **(Fig. 5c)**. These results are consistent with recent chimaera studies ^37,38^ showing that a lack of TET enzyme activity impairs erythropoiesis.

### Tet TKO results in priming defects at enhancers associated with hematopoietic genes

We hypothesised that some of the regulatory elements that are epigenetically perturbed in *Tet* TKO EpiLCs prime lowly expressed developmental genes for future activation **(Fig. 4e)**. In this scenario, these genes would fail to be upregulated upon differentiation of *Tet* TKO cells. To identify such cases, we looked for genes that were not expressed in EpiLC cells or early EBs but significantly lower in *Tet* TKO EBs at day 4, day 6 and day 8 than their WT counterparts (and then remained significantly lower for the rest of the time course). This analysis identified 14, 98 and 297 genes significantly lower than expected at day 4, 6 and 8, respectively **(Fig. 6a, Supplemental Fig. 8a, b)**. Interestingly, the genes that failed to be upregulated in *Tet* TKO EBs also failed to be upregulated in *Dnmt* TKO EBs **(Supplemental Fig. 8c)**.

**Figure 6.**
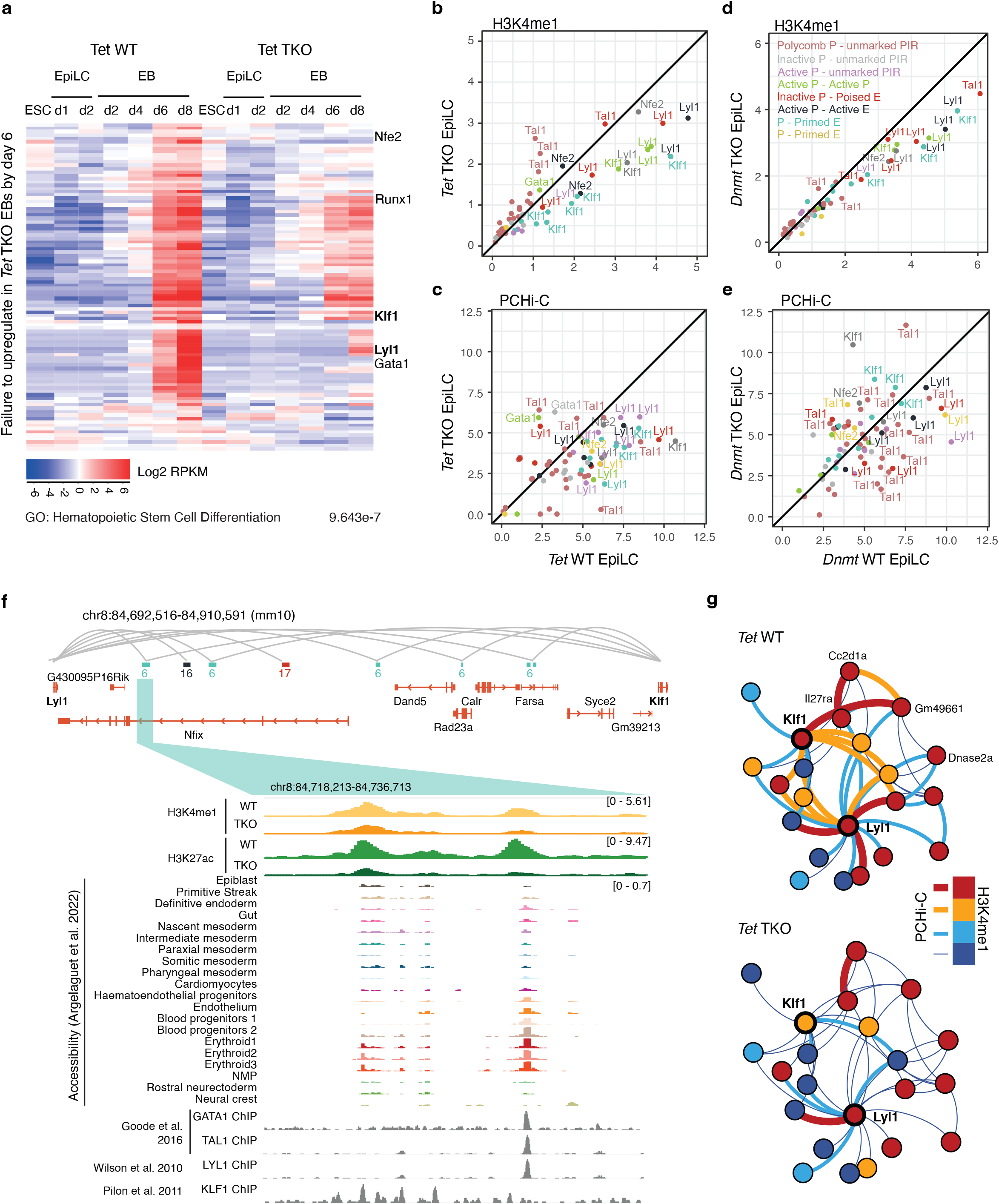
*Tet* or *Dnmt* TKO result in failure to prime enhancers associated with hematopoietic transcription factors **(a)** Heatmap showing genes that are significantly upregulated by day 6 of EB differentiation (relative to ESCs cultured in 2i-LIF) and remain upregulated in day 8 EBs generated from WT cells, but fail to be upregulated in EBs generated from *Tet* TKO cells (according to DESeq and an dynamic fold change filter). Gene ontology analysis identified “Hematopoietic stem cell differentiation” as the top term within which these genes were enriched (below). Key hematopoietic TFs from this term are labelled. **(b-e)** Scatter plots displaying the levels of H3K4me1 (b,d) or the interaction strength by PCHi-C (c, e) in *Tet* TKO (b, c) or *Dnmt* TKO (d, e) EpiLCs relative to respective WT controls at promoter interacting regions (PIRs) involving the indicated blood TFs (labelled in a). H3K4me1 levels are quantified as reads per kilobase of fragment per million (RPKM) whilst PCHi-C values are asinh transformed CHiCAGO scores capped at 3 MAD (median absolute deviation). PIRs are labelled by the gene they interact with and coloured by the cluster that the promoter-PIR pair was assigned to (see Fig. 2b). **(f)** (top) Gene track showing the locations of two blood TF genes, *Lyl1* and *Klf1*, and intervening genes. Grey arcs represent interactions involving *Lyl1* and/or *Klf1* and the location and cluster number of PIRs is indicated and coloured as in Fig. 2b. (Bottom) genome browser image showing the levels of H3K4me1 (orange) and H3K27ac (green) in WT and *Tet* TKO EpiLCs across a primed enhancer region that interacts with *Klf1* and *Lyl1*. Also displayed are public accessibility data from different cell types found in the E8.5 mouse embryo and ChIP data for hematopoietic TFs. **(g)** Subnetwork of *Klf1* and *Lyl1* promoters and their interacting regions (PIRs): Nodes (promoters and PIRs) are coloured by their levels of H3K4me1 (low, medium low, medium high, high). PCHi-C interaction strength between nodes is indicated by colour and line thickness (red = high, blue = low, for thresholding criteria see accompanying material).

Consistent with GO analysis using all differentially expressed genes **(Fig. 5d)**, these lists were significantly enriched for genes associated with hematopoietic stem cell differentiation (adjusted p-value 9.643e-7), and included genes encoding key transcription factors - *Tal1, Nfe2, Runx1, Klf1, Lyl1* and *Gata1* **(Fig. 6a)**. To determine whether the regulatory elements associated with these blood transcription factor genes lose epigenetic priming at the EpiLC stage in the mutants, we plotted the levels of ATAC, H3K4me1 and H3K27ac signal as well as interaction strength at associated PIRs in WT controls versus *Tet* TKO and *Dnmt* TKO EpiLCs **(Fig. 6b-e, Supplemental Fig. 8d-g)**. Strikingly, many of the PIRs that we had previously annotated as active or primed enhancers **(Fig. 2b)** had reduced accessibility, reduced levels of active histone modifications and lowered interaction strength in *Tet* and *Dnmt* TKO EpiLCs, long before the linked genes are expressed **(Fig. 6b-e, Supplemental Fig. 8d-g)**. These data are consistent with a model whereby cyclical turnover of DNA methylation by DNMTs and TETs is important for epigenetic priming of blood enhancers and their interactions with cognate promoters.

This defect was especially prominent for promoter-PIR pairs involving two transcription factor genes: *Klf1* and *Lyl1* **(Fig. 6b-e; Supplemental Fig. 8d-g)**. These are found approximately 200 kb apart in both the mouse (on chromosome 8) and human genomes (on chromosome 19) and they are separated by homologous genes including *Nfix, Dand5*, and *Syce2*, demonstrating evolutionary conservation of the locus **(Fig. 6f, top)**. Interestingly, subnetworks for those two genes revealed that many of these regulatory elements were shared by both *Klf1* and *Lyl1* **(Fig. 6f, g)**. For example, one PIR that interacts with both genes is flanked by H3K4me1 and H3K27ac enriched regions. These histone marks and the strength of the interactions between the elements were substantially reduced upon *Tet* TKO in EpiLC cells **(Fig. 6f, g)**, suggesting loss of epigenetic priming. Next, we analysed published 10x Multi-ome (combined single cell RNA-seq and ATAC-seq) ^62^ data of E8.5 mouse embryos to identify cell type specific accessibility profiles. Strikingly, the *Klf1/Lyl1* interacting regulatory elements that lose epigenetic priming in *Tet* and *Dnmt* TKO EpiLC cells became accessible specifically in hematopoietic progenitor cells and erythroid lineages **(Fig. 6f, bottom)**. Moreover, these accessible elements are later bound by blood associated transcription factors that are not expressed at the EpiLC stage, including GATA1, TAL1 and LYL1, according to reanalysed ChIP-seq data ^63–65^ from hemato-endothelial progenitor cells **(Fig. 6f, bottom)**. Collectively, our data suggest that these regulatory elements function as epigenetically primed enhancers that are required for hematopoietic differentiation. We find that methylation and oxidation by DNMTs and TETs is required to maintain accessibility, active histone modifications and looping interactions between these elements and key blood TF genes.

## Discussion

### Dynamic turnover of DNA methylation at pluripotent stem cell enhancers

Paradoxically, the enzymes that deposit (DNMTs) and remove (TETs) DNA methylation are co-expressed during early mammalian development and target overlapping genomic regions, in particular distal gene regulatory regions such as enhancers ^21,23,24,66,67^. As a consequence, DNA methylation states at gene regulatory elements cycle dynamically in mouse and human pluripotent stem cells, especially during the transition from naïve to formative pluripotency. This raises the possibility that the antagonistic enzymes of the DNA methylation cycle, or the modifications they generate, may be involved in modulating the activity of key gene regulatory elements during early developmental cell fate diversification^22^.

Here, to test this hypothesis, we combine an established *in vitro* cell model of formative pluripotency with genetic mutants that are defective for specific enzymatic steps in the DNA methylation and oxidation cycle. We extensively profile multiple chromatin modifications, DNA methylation, and transcription, and we developed a novel clustering approach to track promoter-gene regulatory element interactions during mouse pluripotency state transitions. We believe that this rich resource, containing more than 300 new datasets, will be of great value to the community. We find that during the exit from pluripotency, dynamic turnover of cytosine methylation is required to activate enhancers that act in mouse pluripotent stem cells. Further, the cycle of DNA methylation and oxidation is also necessary to prime enhancers for future activation, and we show that aberrant enhancer priming in TET and DNMT mutants results in the failure to upregulate key transcription factor genes within the hematopoietic lineage tree. Our data thus suggest that the role of DNA oxidation at gene regulatory regions goes beyond removing DNA methylation as a repressive epigenetic mark, to keep the local chromatin accessible for trans-acting factors that mediate enhancer function ^68,69^. We propose that in addition to this role, enhancer DNA demethylation functions in a dynamic DNA methylation and oxidation cycle to establish an active and primed enhancer landscape that controls cell-type specific gene expression programmes in pluripotent stem cells and their differentiated progeny.

### Potential mechanisms of enhancer priming by cyclical DNA methylation and oxidation

Enhancers have been grouped into functional categories based on post-translational histone modifications they harbour, including primed (H3K4me1), active (H3K4me1 and H3K27ac) and poised (H3K4me1 and H3K27me3; ^41,42^). These post-translational histone modifications are deposited at enhancers by MLL3/MLL4 (H3K4me1), P300/CBP (H3K27ac), and Polycomb repressive complex 2 (H3K27me3). We demonstrate that interfering with DNA methylation dynamics affects the chromatin states at all three enhancer classes, and that the lack of DNMT activity causes changes in enhancer chromatin signatures that are distinct from those resulting from the absence of catalytically active TETs (**Fig. 7**). H3K4me1 occupancy at regulatory elements is markedly increased during the ESC to EpiLC transition, which we found is dependent on both DNMT and TET activity. In contrast, H3K27 acetylation is depleted at enhancers in oxidation deficient cells, but persists in the absence of DNMT activity. The Polycomb mark H3K27me3 is broadly redistributed in *Tet* knockout EpiLC, resulting in a pronounced gain of H3K27me3 at poised enhancers. Our data suggest that oxidation derivatives of 5mC (5hmC, 5fC or 5caC) may be important for H3K4me1 deposition at enhancers. Intriguingly, we have previously identified WDR5, a member of the WRAD complex that may be involved in targeting histone methyltransferases to specific genomic sites ^70^, as strongly interacting with 5fC modified DNA ^71^. This interaction may provide a mechanistic link between oxidation derivatives and MLL3/MLL4 mediated H3K4me1 deposition at enhancers **(Fig. 7)**.

**Figure 7.**
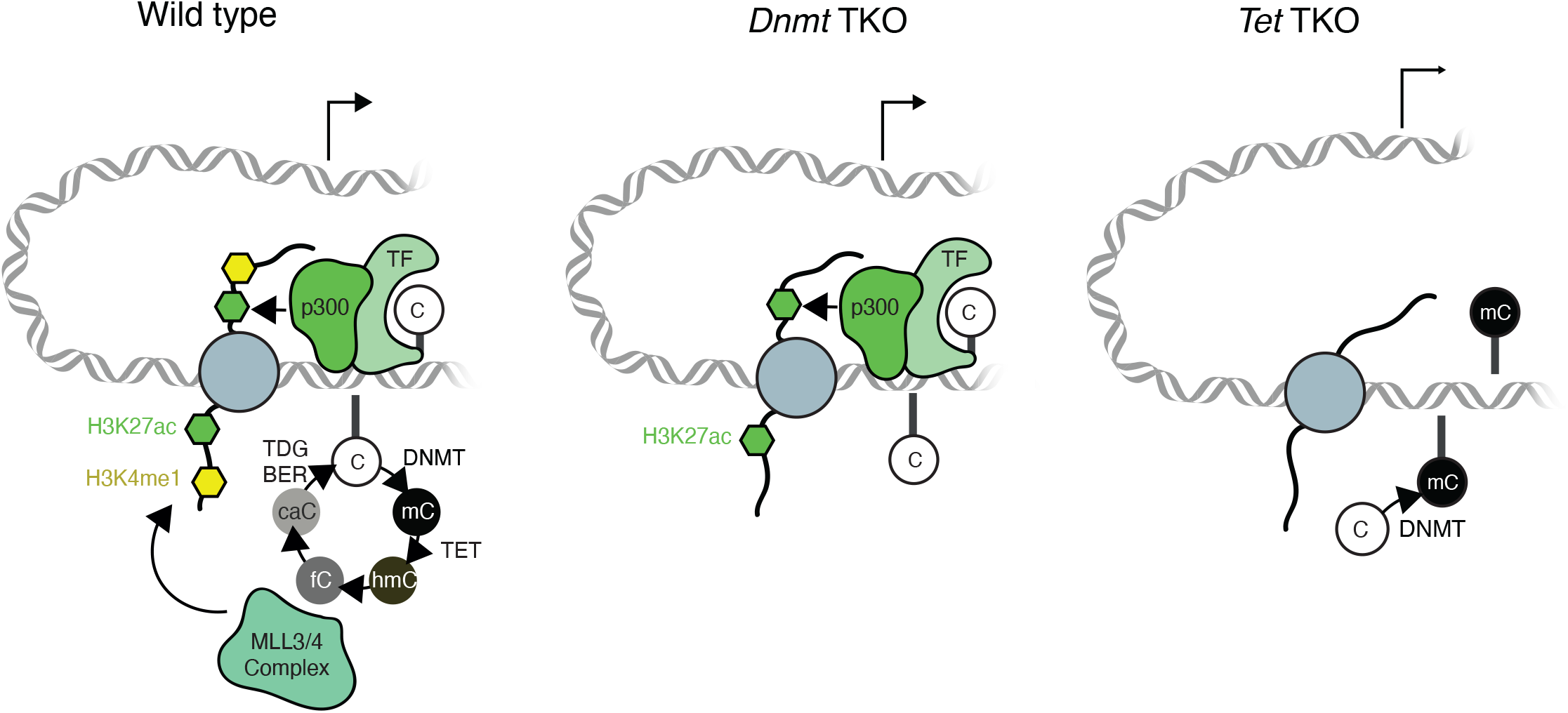
Cyclical DNA methylation turnover may be required for epigenetic priming of enhancers Model figure describing the epigenetic landscape of an active gene in EpiLCs. H3K4me1 deposition is perturbed in both *Dnmt* and *Tet* TKO cells, suggesting that both enzymatic activities and DNA methylation turnover are required. MLL3/4, part of the machinery that deposits H3K4me1, is depicted as binding 5fC as an example of how methylation turnover could be important. H3K27ac deposition is perturbed only in *Tet* TKO cells, suggesting that this mark requires unmethylated cytosine. We speculate here that methylation sensitive transcription factors recruit p300 to deposit this H3K27ac, and that this would not be possible in *Tet* TKO cells where 5mC accumulates. At an epigenetically primed enhancer that regulates a developmental gene, such as a gene encoding a hematopoietic transcription factor, cyclical DNA methylation turnover in the E5.5 epiblast by TET and DNMT enzymes produces 5mC and oxidative derivatives. These derivatives recruit MLL3/4 containing complexes that in turn deposit H3K4me1 at primed enhancers, facilitating three-dimensional interactions with cognate promoters and forming an epigenetic state permissive for enhancer and gene activation following differentiation (top, WT). In *Tet* or *Dnmt* TKO cells DNA methylation turnover is perturbed - oxidative derivatives are not produced - and as such H3K4me1 can not be targeted efficiently. This diminished priming prevents proper gene activation upon differentiation.

Our data further imply that H3K27ac can persist when H3K4me1 is depleted, in agreement with a recent study reporting that H3K27ac is gained at hundreds of enhancers independently of MLL3/4 activity or H3K4me1 binding during ESC differentiation ^72^. The requirement for enhancer priming, and the underlying mechanisms, are likely to be developmental context dependent. Ectoderm-specifying enhancers, unlike endoderm- and mesoderm-associated enhancers, are hypomethylated and accessible long before ectoderm specification during mouse gastrulation ^10^. However, the low levels of DNA methylation at these enhancers are TET-dependent. It will be interesting to determine whether a dynamic DNA methylation turnover occurs at ectoderm enhancers during early mouse development, and if it does, how the kinetics differ compared to endo- and mesoderm specifying enhancers. Moreover, we identify here a subset of mesodermal enhancers that are epigenetically primed by H3K4me1 in epiblast-like cells, and so early epigenetic priming is not completely exclusive to ectoderm enhancers.

A recent study demonstrated that H3K27 acetylation is dispensable for gene activation during the ESC to EpiLC cell fate transition ^73^. One possible explanation is that substrates other than H3K27, including non-histone targets such as transcription factors, are the functionally relevant targets of P300/CBP’s lysine acetyltransferase activity at enhancers ^74,75^. We note that our model **(Fig. 7)** is entirely compatible with this scenario, as it places the DNA methylation turnover upstream of enhancer-associated chromatin modifiers and their histone and non-histone targets.

### Rewiring of 3D promoter chromatin interactions upon perturbed DNA methylation turnover

Intriguingly, we found that in DNA methylation or oxidation deficient cells, regulatory interactions are disrupted and active chromatin marks are reduced at specific promoter interacting regions, long before the linked target genes are expressed during development. This priming defect appears to be an enhancer-specific rather than a global effect, with genes encoding key blood lineage transcription factors including *Klf1* and *Lyl1* among those affected. Catalytic inactivation of MLL3/4 has been shown to result in a loss of H3K4me1 at enhancers ^76^, and MLL3/4 and H3K4me1 occupancy at enhancers correlates with increased levels of chromatin interactions via recruitment of cohesin ^77^. Interestingly, MLL3/4 activity is specifically required at enhancers that gain H3K4me1 during ESC differentiation, and enhancer-promoter contacts at these de novo enhancers are severely disrupted in the absence of MLL3/4 ^78^. This indicates that MLL3/4 may be necessary to maintain enhancers in a plastic state that allows them to respond efficiently to developmental cues. How the dynamic turnover of DNA methylation contributes to enhancer plasticity, and which role it plays at MLL3/4-dependent versus -independent enhancers ^78^ warrants future investigation.

### Aberrant DNA methylation in haematological malignancies

Finally, our data may also provide an explanation for the paradoxical finding that mutations in *DNMT* and *TET* genes both have been causally linked to a range of blood cancers. For example, *TET2* mutations are present in approximately 50% of chronic myelomonocytic leukaemia (CML) and 10% of acute myeloid leukaemia (AML) cases ^33^. Mutations in *DNMT3A* were first reported in AML patients and have since been discovered in several other haematological malignancies, pointing to DNMT3A as an important tumour suppressor ^79^. We propose that the antagonistic activities of DNMTs and TETs converge on the dynamic methylation turnover at distal regulatory elements, which is required for their activity. The dysregulation of DNA methylation and oxidation at enhancers may contribute to the aetiology and pathology of blood cancers ^80^, which could inform novel diagnosis approaches and treatment strategies to improve patient care.

## Data Availability

Sequencing data will be available in the NIH GEO database upon publication. Quantitated data for interaction pairs (which forms the basis for clustering, UMAP and the majority of plots) will be provided as a data table under the same accession. Further accompanying material including analysis details are available in a GitHub repository (https://github.com/ChristelKrueger/Exit-from-Pluripotency) and organised in a website at https://christelkrueger.github.io/Exit-from-Pluripotency/.

## Acknowledgements

The authors would like to thank all members of the Reik lab for helpful discussions. We especially thank Stephen Clark and Ricard Argelaguet for discussions and ideas. The authors would like to thank the Bioinformatics Facility (particularly Simon Andrews and Felix Krueger) at the Babraham Institute for helpful discussion and for processing sequencing data. We also thank the Proteomics facility (particularly David Oxley and Judith Webster) for performing DNA mass spectrometry experiments. A.P. was funded by a Sir Henry Wellcome Fellowship (215912/Z/19/Z). C.K., S.S. and S.W. were funded by the Babraham Institute’s UKRI-BBSRC core capability grant (BBS/E/B000X0000) and the Epigenetics Institute Strategic Programme (BBS/E/B/000C0425). T.L. was funded by the Wellcome Trust 4-Year PhD Programme in Stem Cell Biology and Medicine and the University of Cambridge, UK (203813/Z/16/A and 203813/Z/16/Z). S.S. was supported by a UKRI MRC Rutherford Fund Fellowship (MR/T016787/1) and a Career Progression Fellowship from the Babraham Institute. W.R. was funded by the Epigenetics Institute Strategic Programme and Wellcome Investigator (210754/B/18/Z) and Collaborative Awards (220379/Z/20/Z).

## Contributions

A.P. and C.K. designed the study with contributions from T.L., S.S. and W.R. A.P. cultured cells, performed experiments, collected and analysed data and wrote the manuscript. C.K. analysed data, developed the novel clustering approach and visualisation methods presented and wrote the manuscript. T.L. cultured cells, performed experiments and collected data. S.W. processed PCHi-C data. S.S. assisted with PCHi-C experiments, supervised the study and wrote the manuscript. W.R. supervised the study and wrote the manuscript.

## Methods

### Experimental methods

#### Cell culture

Naïve mESCs were cultured in 2i-LIF conditions ^81^ consisting of N2B27 media (N2 & B27; Life Technologies) supplemented with 10 ng/mL mLIF (Cambridge Stem Cell Institute [SCI]), 1 mM MEK inhibitor PD0325901 (SCI), and 3 mM GSK3 inhibitor CHIR99021 (SCI) on gelatin coated plates. EpiLC transition was induced by plating 1×10^3^ naive mESCs on a human plasma fibronectin (Millipore, FC010) coated plate in N2B27 medium supplemented with 20 ng/mL activin A (SCI), 12 ng/mL bFGF (SCI), and 1% KSR (Gibco) ^4^. The media was changed after 24 hours and cells harvested after 48 hours.

*Dnmt* TKO mESCs were a generous gift from Dirk Schübeler ^53^. *Tet* TKO ESCs were a generous gift from Guo-Liang Xu ^28^. Cells were tested for mycoplasma contamination using a MycoAlert Mycoplasma Detection Kit (Lonza, LT07-318) and found to be negative.

#### RNA-seq

RNA was isolated from cells using QIAshredder columns and RNeasy Mini kits (QIAGEN) according to the manufacturer’s instructions. Isolated RNA samples were treated with TURBO DNase (Thermo Fisher Scientific, AM2238) before samples were submitted for library preparation and sequencing at the The Wellcome Trust Sanger Institute.

#### ChIP-seq and HMCP-seq

ChIP-seq libraries were prepared as previously described ^82^. Chromatin from 10 million cells was sonicated to between 200-600 bp using a Bioruptor Plus (Diagenode). 100µL of Dynabeads Protein G (Invitrogen, 10003D) were used per immunoprecipitation. Libraries were prepared using the NEBNext Ultra II Library Prep Kit for Illumina (New England Biolabs, E7645) following the manufacturer’s instructions. Libraries were size selected after amplification using Agencourt Ampure XP beads (Fisher Scientific, 10136224) prior to pooling and sequencing.

The following antibodies were used: H3K4me3 (Abcam, ab8580), H3K27me3 (Millipore, 07-449), H3K4me1 (Active Motif, 39297), H3K27ac (Abcam, ab4729), CTCF (Millipore, 07-729).

HMCP-seq, a method for 5hmC pulldown, was performed using a kit (provided by Cambridge Epigenetix) according to manufacturer’s instructions without modification.

#### ATAC-seq

ATAC-seq was performed based on published protocols ^83^ using reagents from a Nextera sequencing kit (Illumina, FC-121-1030). Cells were detached using TrypLE Express (Gibco, 12604013), counted, and 10,000 cells aliquoted into DNA LoBind tubes (Eppendorf, 0030108051). Cells were pelleted for 5 minutes at 500g and 4ºC before lysis in 10µL of cold lysis buffer (10mM Tris HCl pH7.4, 10mM NaCl, 3mM MgCl2, 0.1% IgePal CA-630) and immediate repeated centrifugation. Supernatant was removed and the nuclear pellet was suspended in 20µL of transposition mix (1x TD buffer and 0.5µL TDE1 TN5 enzyme) before incubation at 37ºC for 30 minutes with mixing at 1000 RPM using a heated shaker. Samples were purified immediately using a MinElute DNA purification kit (QIAGEN). Transposed DNA was amplified using the adapters described by Buenrostro et al., 2015 at 1.25µM and NEBNext High Fidelity 2x Mastermix (New England Biolabs, M0541) in a 50µL reaction (5 minutes at 72ºC, 30 seconds at 98ºC, followed by 13 cycles of 10 seconds at 98°C, 75 seconds at 65°C and 60 seconds at 72°C). Samples were purified and size selected using Agencourt Ampure XP beads(Fisher Scientific, 10136224) prior to pooling and sequencing.

#### WGBS

As an input for WGBS 400 ng of genomic DNA was fragmented to between 300 and 400 bp using a Covaris E220 water bath sonicator following manufacturer’s instructions. The DNA ends were repaired using NEBNext Ultra II Library Prep (New England Biolabs, E7645) reagents and methylated adapters were annealed according to the manufacturer’s instructions. Fragments were purified using Agencourt Ampure XP beads (Fisher Scientific, 10136224) and bisulfite converted using a Zymo EZ Methylation Gold kit (Zymo, D5005) according to the manufacturer’s instructions and eluting the converted DNA in 15µL of provided elution buffer. Bisulfite treated DNA was amplified using NEBNext index primers (NEB, E7335) and KAPA HiFi Uracil+ mastermix (Roche, KK2801/2) using the following protocol: 1 minute at 98ºC, 10 cycles of 15 seconds at 98ºC, 30 seconds at 65ºC, 30 seconds at 72ºC followed by a final extension for 1 minute at 72ºC. Libraries purified using 0.8x volumes of Agencourt Ampure XP beads (Fisher Scientific, 10136224) to remove contaminating adapters.

#### PCHi-C

PCHi-C was performed as described in and ^45^ using in-nucleus ligation ^84^. The SureSelect Target Enrichment system (Agilent Technologies) described by ^45^.

#### DNA Mass Spectrometry

A total of 200 ng of genomic DNA was digested using 5U of DNA Degradase Plus (Zymo Research, E2020) overnight at 37ºC and the base content quantified using a nanoHPLC and Q Exactive mass spectrometer (as described in ^85^).

### Processing of sequencing data

#### RNA-seq

RNA-seq libraries were sequenced as 75 base paired-end runs on an Illumina HiSeq2500 instrument. Quality and adapter trimming were performed using Trim Galore v0.6.1 (using Cutadapt v1.18) ^86^. Mapping to the mouse reference genome (GRCm38) was performed using Hisat2 v2.1.0 (--no-softclip) ^87^ guided by known splice sites from Ensembl annotation v90. Differential gene expression was determined using DESeq2 and the dynamic fold-change filter within Seqmonk V1.48.1 ^88,89^. Gene ontology analysis was performed using Enrichr ^90–92^.

#### ATAC-seq, ChIP-seq and HMCP-seq

ATAC-seq libraries were sequenced as 50 base single-end runs on an Illumina HiSeq2500 instrument. ChIP-seq libraries were sequenced as 50 base paired-end runs on an Illumina NovaSeq instrument. HMCP-seq libraries were sequenced as 75 base single-end runs on an Illumina NextSeq500 instrument. Quality and adapter trimming were performed using Trim Galore v0.6.5 (using Cutadapt v2.3). Mapping to the mouse reference genome (GRCm38) was performed using Bowtie2 v2.4.1 ^93^.

#### WGBS

WGBS-seq libraries were sequenced as 150 base paired-end runs on an Illumina HiSeq2500 instrument. Quality and adapter trimming were performed using Trim Galore v0.6.6 (using Cutadapt v2.3). Bismark v0.23.0 ^94^ was used to align DNA reads to the bisulfite converted GRCm38 mouse genome then perform methylation calling.

#### PCHiC

PCHiC-seq libraries were sequenced as 75 base paired-end runs on an Illumina HiSeq2500 instrument. Quality and adapter trimming were performed using Trim Galore v0.6.5 (using Cutadapt v2.3). Mapping to the mouse reference genome (GRCm38) was performed using the HiCUP pipeline v0.7.2 ^95^.

### Quality Control

Details regarding data analysis can be found in the accompanying material (https://christelkrueger.github.io/Exit-from-Pluripotency/). Briefly, for quality control, trimming, mapping, FastQC ^96^ and FastQScreen ^97^ reports were collated using MultiQC ^98^. WGBS and PCHiC sample quality was assessed with Bismark and HiCUP reports, respectively. Post-mapping QC was performed with Seqmonk and data distribution, grouping, duplication, coverage and enrichment was assessed. Coverage outliers and regions represented differently between TET and DNMT lines were excluded from further analysis.

### Quantitation and Data Integration

PCHiC data is given as linear CHiCAGO ^99^ scores; if an interaction was found in both directions, the maximum score was used. To integrate PCHiC with ChIP, ATAC and HMCP data, reads were quantitated across HindIII fragments and normalised for length and total number of reads in the sample (RPKM). WGBS data was quantitated across 100 CpG using the Seqmonk methylation quantitation pipeline (min count per position = 1, min number of observations = 20) and is expressed in percent. To obtain methylation levels on HindIII fragment resolution, fragments were overlapped with 100 CpG windows. If a HindIII fragment overlapped with more than one 100 CpG window, the average was taken. All values are replicate averages.

PCHiC interacting pairs form the basis for UMAP and network representations. Quantitated data from different modalities were collated in a data table in which rows represent an interaction pair and columns are attributes of the interaction (PCHiC) or the respective bait and PIR (ATAC, ChIP, HMCP, WGBS). Values are linear. For some visualisations (mainly were level of the mark was represented as colour) capped values were used to obtain a more reasonable scale. Caps were introduced at 3x MAD (mean absolute deviation). Both bait and PIR were annotated with promoters (overlapping gene start coordinates); if more than one promoter was present in the HindIII fragment, the most upstream was used. Data tables for both raw and capped quantitations by interaction pair are provided as processed data on GEO.

### UMAP for interaction pairs, clustering and network

To generate a two dimensional representation of the interaction pair attributes, Uniform Manifold Approximation and Projection for Dimension Reduction (UMAP) was performed. Specifically, centred and scaled linear quantitations of the different sequencing data modalities from the EpiLC TET WT condition were subjected to UMAP using the uwot package ^100^ with spectral initialisation (Settings: min_dist = 0.01, n_neibors = 2, scale = TRUE). Clustering was performed on a 30 dimensional projection of the same input data by Expectation Maximisation for 20 clusters using WEKA ^101^.

Global interactions were also represented as a Canvas ^46^ style network. The R igraph package ^102^ was used to construct an undirected graph in which genomic regions (HindIII fragments) are represented as vertices/nodes and interactions are represented as edges. A combined network was produced using all significant interactions in any of the conditions. The network was visualised with a force-directed layout (ForceAtlas2) in Gephi v0.9 ^103^. This representation pulls highly interacting regions closer together while less interacting regions are kept apart. Details can be found in the accompanying material (https://christelkrueger.github.io/Exit-from-Pluripotency/).

## Figure Legends

**Supplemental Figure 1 (related to Fig.1).**
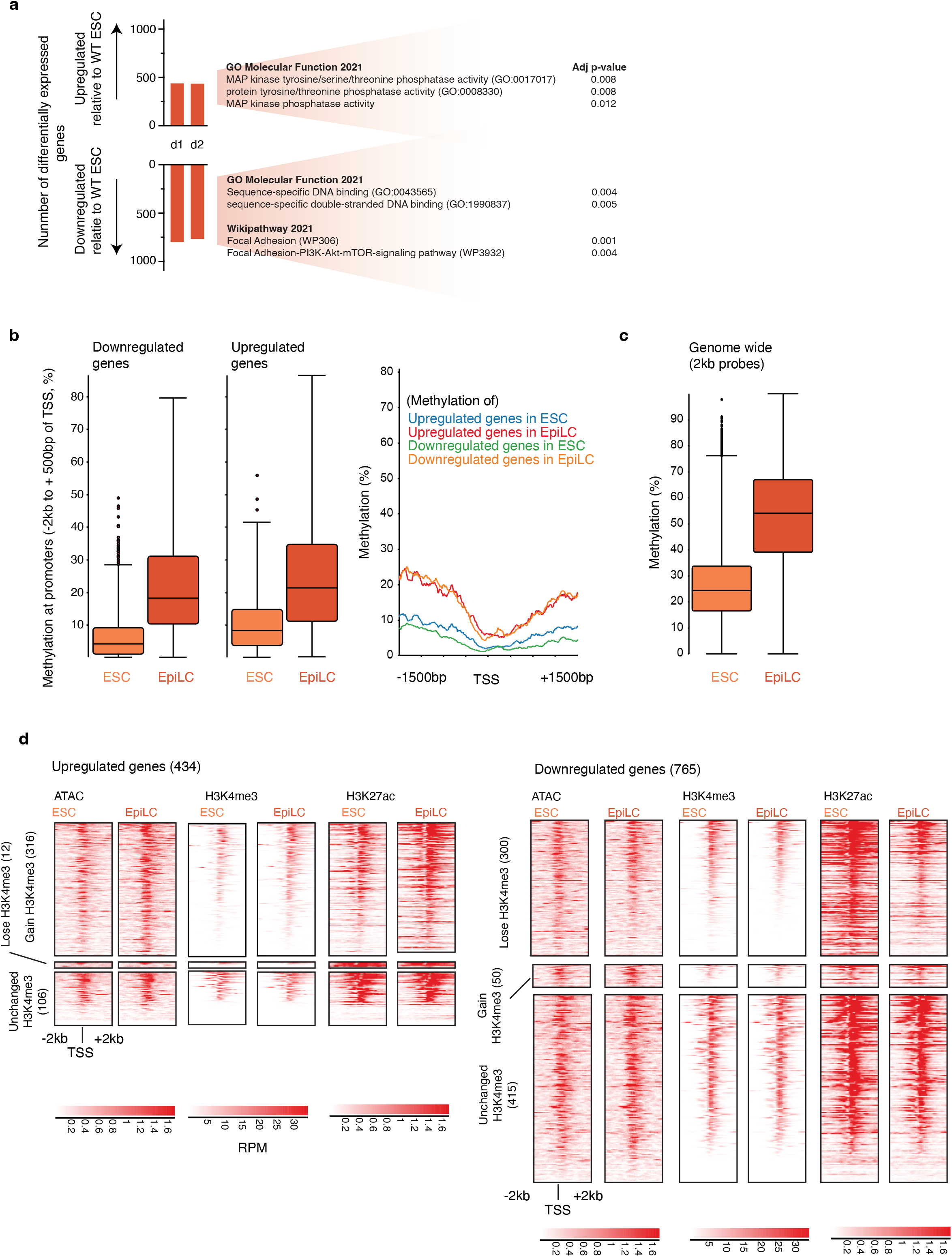
**(a)** Number of differentially expressed genes by mRNA-seq (DESeq and dynamic fold change filter) upon ESC to EpiLC transition after 1 or 2 days (left). GO term enrichment and adjusted p-values associated with these terms for the upregulated and downregulated genes are displayed (right). **(b)** (Left) Box and whisker plots showing CpG methylation (%) across the promoters (−2000 and +500 bp of transcriptional start site, TSS) of genes upregulated and downregulated following two days of ESC to EpiLC transition. (Right) Line plots showing average CpG methylation (% of CpGs) in regions +/- 1500 bp of genes upregulated and downregulated following two days of ESC to EpiLC transition. Levels are shown for ESCs and EpiLCs. **(c)** Box and whisker plots showing CpG methylation genome wide, where tiled 2kb probes have been quantified, in ESCs and EpiLCs. **(d)** Aligned probe plots showing ATAC-, H3K4me3 ChIP- and H3K27ac ChIP-seq data across 4kb regions centred at the TSS of genes upregulated (left) and downregulated (right) upon ESC to EpiLC transition. Promoters are split based on whether H3K4me3 accumulates, is depleted or is unchanged across these regions upon ESC to EpiLC transition (significant according to DESeq, adjusted p-value < 0.05).

**Supplemental Figure 2 (related to Fig. 1).**
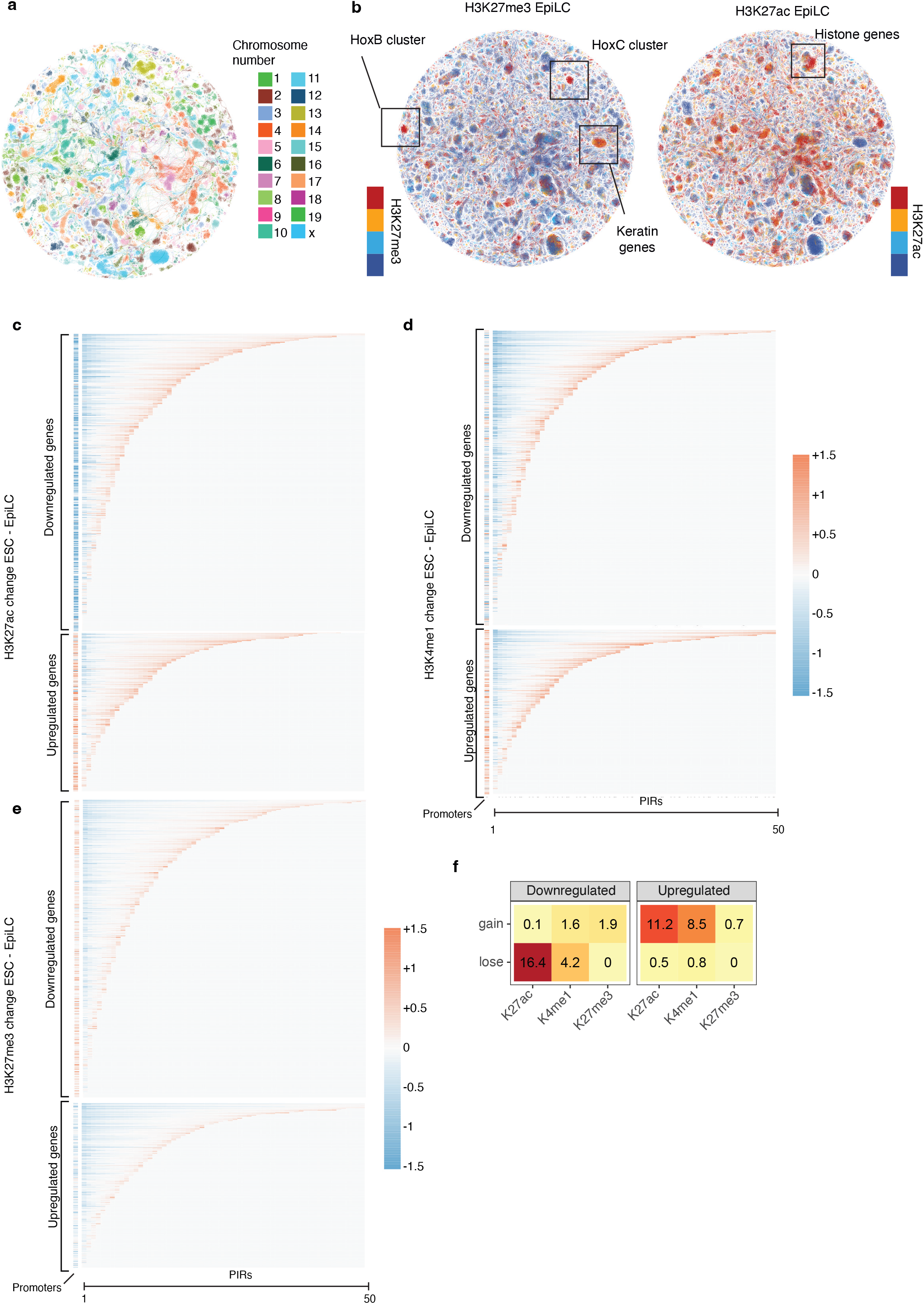
**(a, b)** Force directed graph of promoters and promoter interacting regions (PIRs)(nodes) connected by the interactions between them (edges) generated using Promoter Capture Hi-C (PCHi-C) data (similar to Canvas in ^46^)). Edges are coloured by **(a)** chromosome or **(b)** levels of H3K27me3 and H3K27ac (red = high, blue = low, for thresholding criteria see accompanying material). The location of genes known to be repressed (HoxB, HoxC, Keratin genes) or active (Histone genes) in EpiLCs are labelled. **(c-e)** heatmaps showing change in H3K27ac (c), H3K4me1 (d) and H3K27me3 (e) levels upon ESC to EpiLC transition (RPKM difference) across the promoters and PIRs of genes that are significantly upregulated and downregulated upon ESC to EpiLC transition at the level of mRNA. The top 50 PIRs that interact with each promoter are displayed, and these are sorted left to right based on extent of histone modification change (loss to gain). **(f)** A summary of the data shown in (c-e). The percentage of PIRs associated with transcriptionally downregulated (left) or upregulated (right) genes that gain or lose each histone modification (RPKM change of >1).

**Supplemental Figure 3 (related to Fig. 1).**
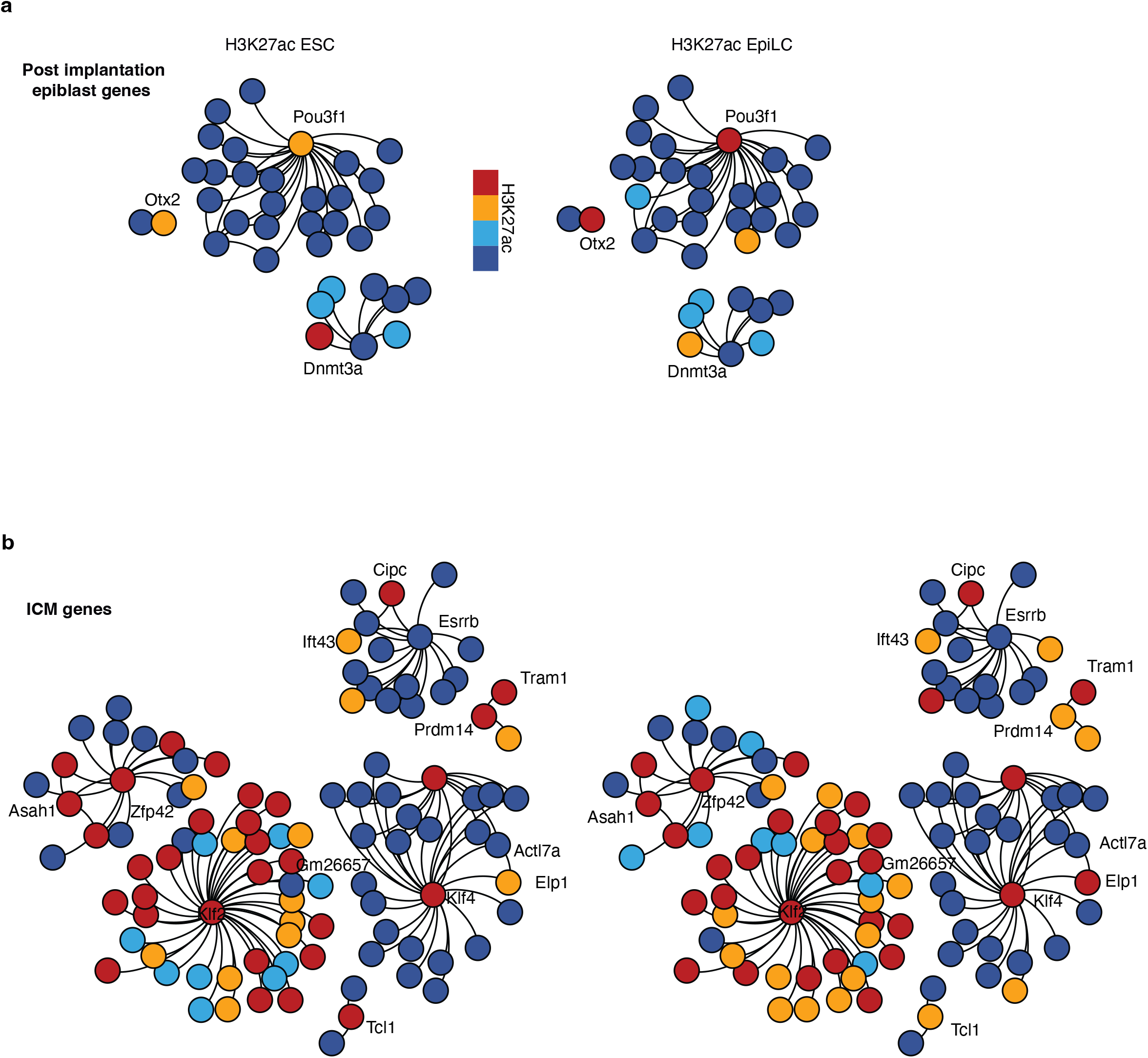
**(a, b)** Force directed graph of promoters and promoter interacting regions (PIRs) (nodes) connected by the interactions between them (edges) generated using Promoter Capture Hi-C (PCHi-C) data. Subclusters are centred on genes of interest and their PIRs coloured by the levels of H3K27ac (low, medium low, medium high, high) in ESCs (left) and EpiLCs (right).

**Supplemental Figure 4 (related to Fig. 2).**
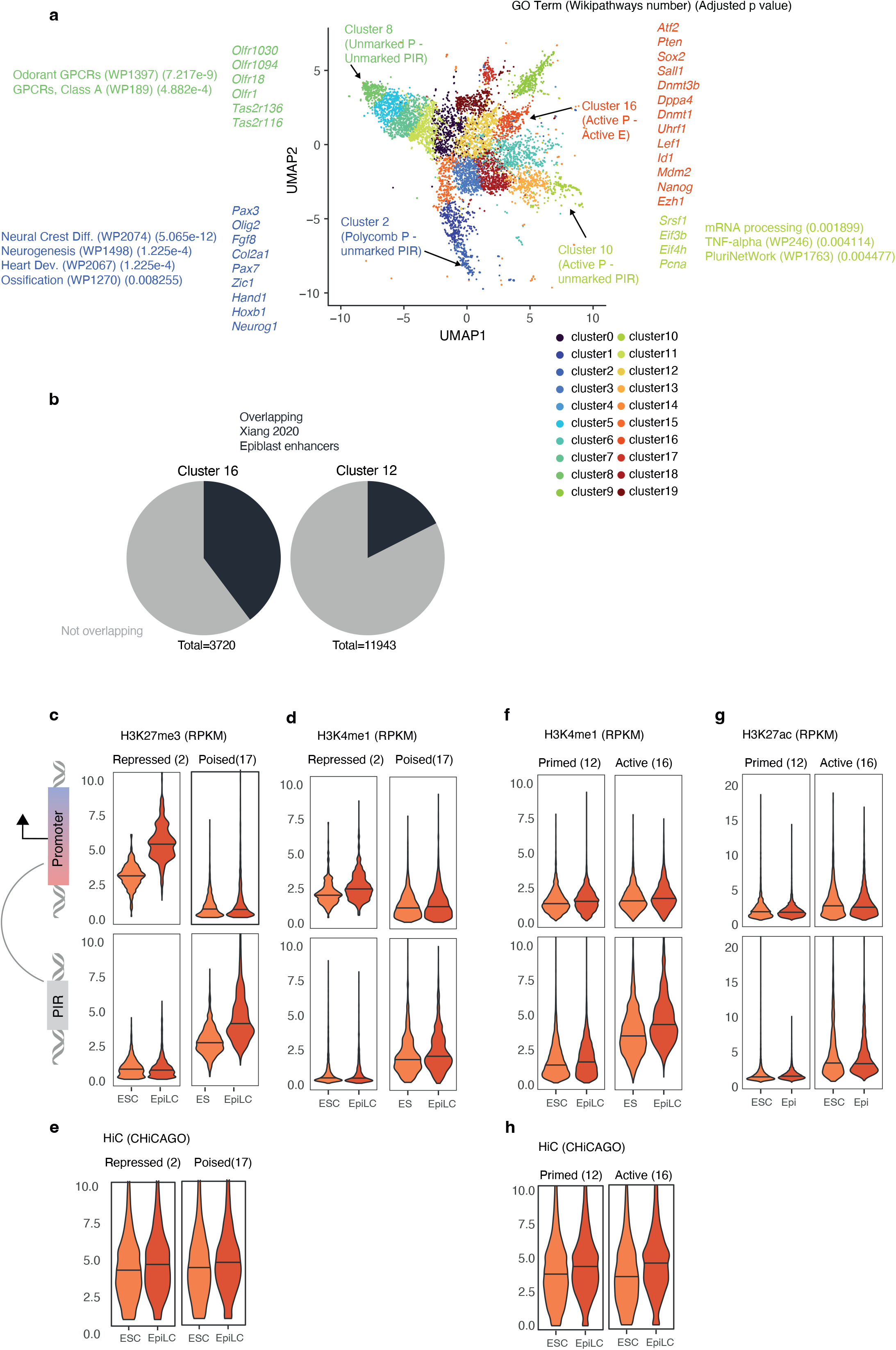
**(a)** Result from the computational approach described in (Fig. 2a) plotted using uniform manifold approximation and projection (UMAP), coloured by clusters generated through expectation maximisation based clustering. Some key clusters are labelled based on interrogation of the histone modifications found at the promoters and PIRs. P = promoter, E = enhancer. Example genes that are associated with the promoters found in clusters 2, 8, 10 and 16 are shown as well as gene ontology categories that are enriched for genes within those clusters and associated adjusted p-values illustrating significance of these enrichments. **(b)** The fraction of PIRs from clusters 16 and 12 that overlap with epiblast enhancers defined by Xiang et al. ^51^. **(c-e)** Violin plots displaying levels of H3K27me3 (c), H3K4me1 (d) and interaction scores (e) in ESC and EpiLC cells across promoters (top) and PIRs (bottom) in clusters 2 and 17 - pairs consisting of polycomb repressed promoters and poised enhancers, respectively. **(f-h)** Violin plots displaying levels of H3K4me1 (f), H3K27ac (g) and interaction scores (h) in ESC and EpiLC cells across promoters and PIRs in clusters 12 and 16 - pairs consisting of primed and active enhancers, respectively. (c-h) ChIP data are quantified as reads per kilobase of fragment per million (RPKM) and interaction data are asinh transformed CHiCAGO scores capped at 3 MAD.

**Supplemental Figure 5 (related to Fig. 3).**
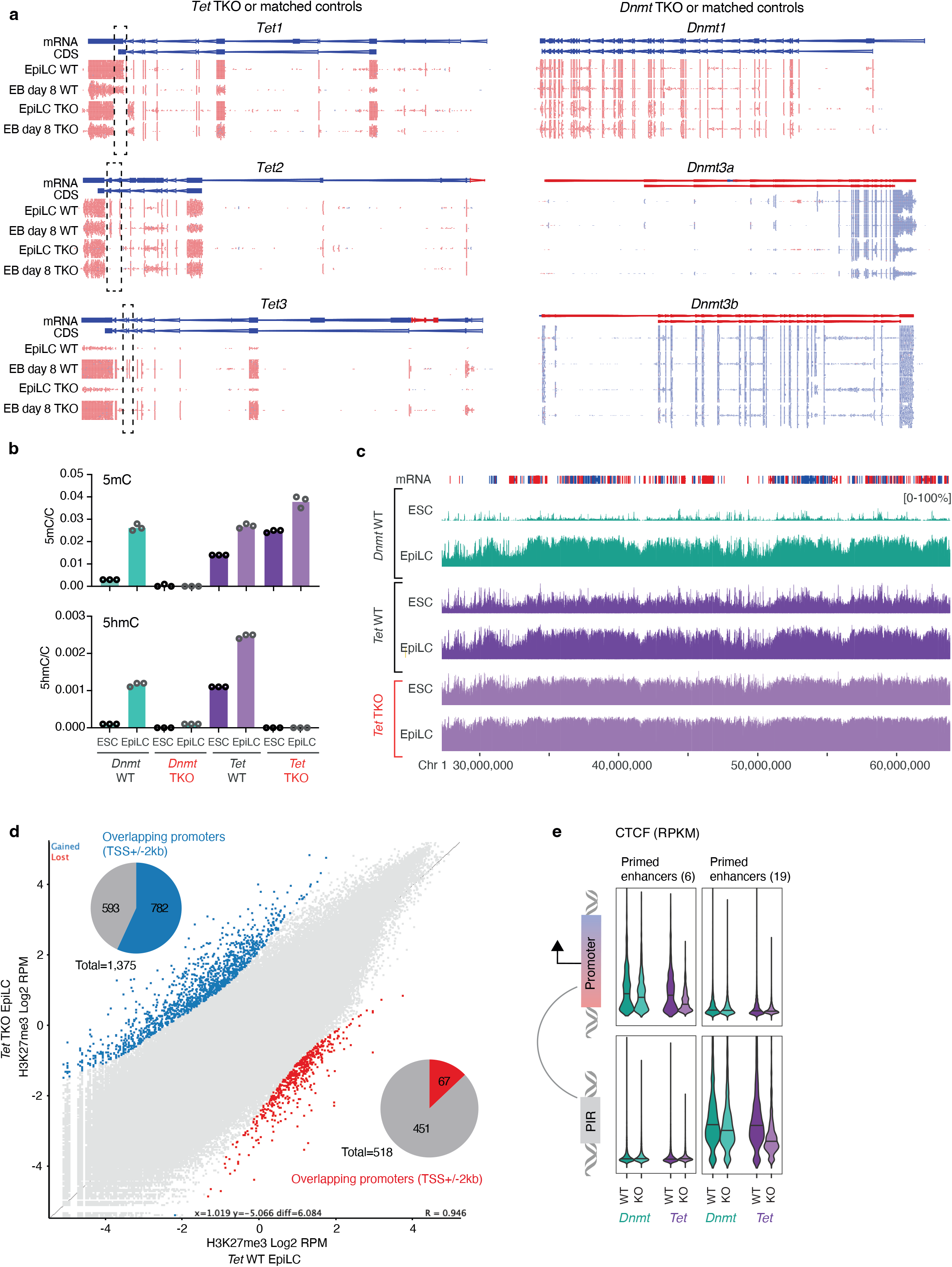
**(a)** Genome browser images showing aligned RNA-seq data from WT and *Tet* TKO cells (left) or *Dnmt* TKO cells (right) grown as EpiLCs or differentiated to embryoid bodies (EBs) for 8 days. The location of mRNA transcripts and the coding DNA sequence (CDS) are displayed. The catalytic domains removed from *Tet1, 2* and *3* in *Tet* TKO cells are indicated by dashed boxes. **(b)** Quantity of 5mC (top) and 5hmC (bottom) as a proportion of total C in WT, *Tet* (purple) and *Dnmt* (green) TKO ESCs (2i-LIF) and EpiLCs. **(c)** Genome browser image showing whole genome bisulfite data (WGBS) across *Dnmt* WT, *Tet* WT and *Tet* TKO cells. The percentage of cytosine methylation at CpG dinucleotides is plotted. **(d)** Scatter plot of H3K27me3 levels (Log_2_ RPM) over 2kb genome-wide tiles in *Tet* WT vs TKO EpiLCs. Regions that significantly gain H3K27me3 (blue) and lose K27me3 (red) are highlighted (intensity difference filter within Seqmonk including multiple testing correction). The fraction of each category overlapping a promoter (transcriptional start site [TSS] +/- 2kb) is plotted. **(e)** Violin plots displaying levels of CTCF (RPKM) across promoters (top) and PIRs (bottom) of pairs in clusters 6 and 19 (involving promoters interacting with primed enhancers).

**Supplemental Figure 6 (related to Fig. 4).**
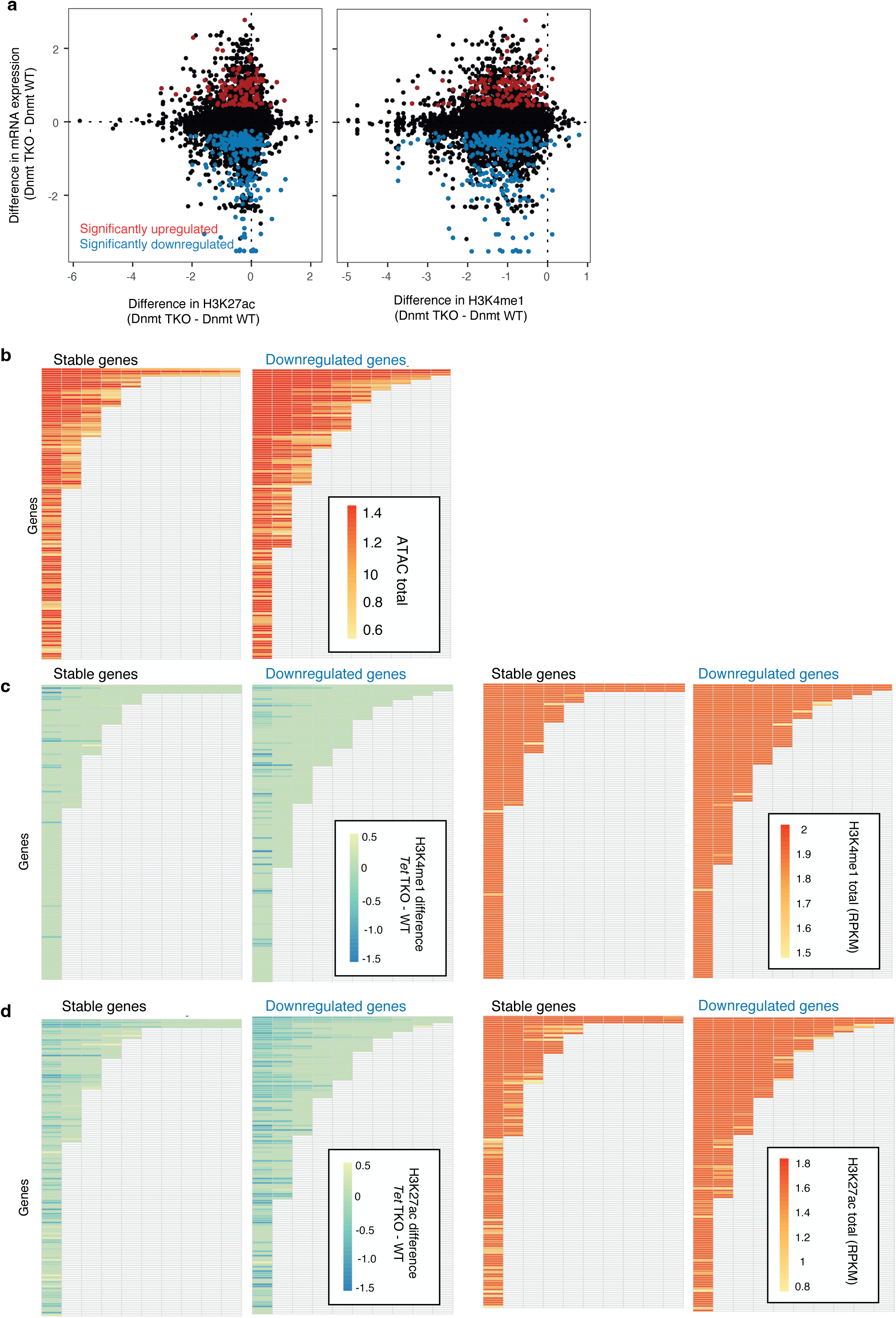
**(a)** Change in mRNA expression against change in H3K27ac and H3K4me1 levels in *Dnmt* TKO cells relative to matched WT controls across active enhancers (defined in Fig. 2). Enhancers linked to differentially expressed genes are indicated. **(b)** Heatmap showing the levels of accessibility (ATAC-seq signal) in WT EpiLCs at enhancer elements linked to genes that are downregulated or stably expressed upon *Tet* TKO EpiLCs. Rows represent genes and columns represent enhancers linked to those genes. Genes are sorted vertically by enhancer number. **(c, d)** Heatmaps showing the change in H3K4me1 (c) and H3K27ac (d) levels at enhancer elements linked stable genes or genes downregulated in *Tet* TKO cells (left). The same heatmaps are plotted on the right but coloured by the starting levels of H3K4me1 (c) and H3K27ac (d) in WT EpiLCs. Rows represent genes and columns represent enhancers. Genes are sorted by the number of interacting enhancers.

**Supplemental Figure 7 (related to Fig. 5).**
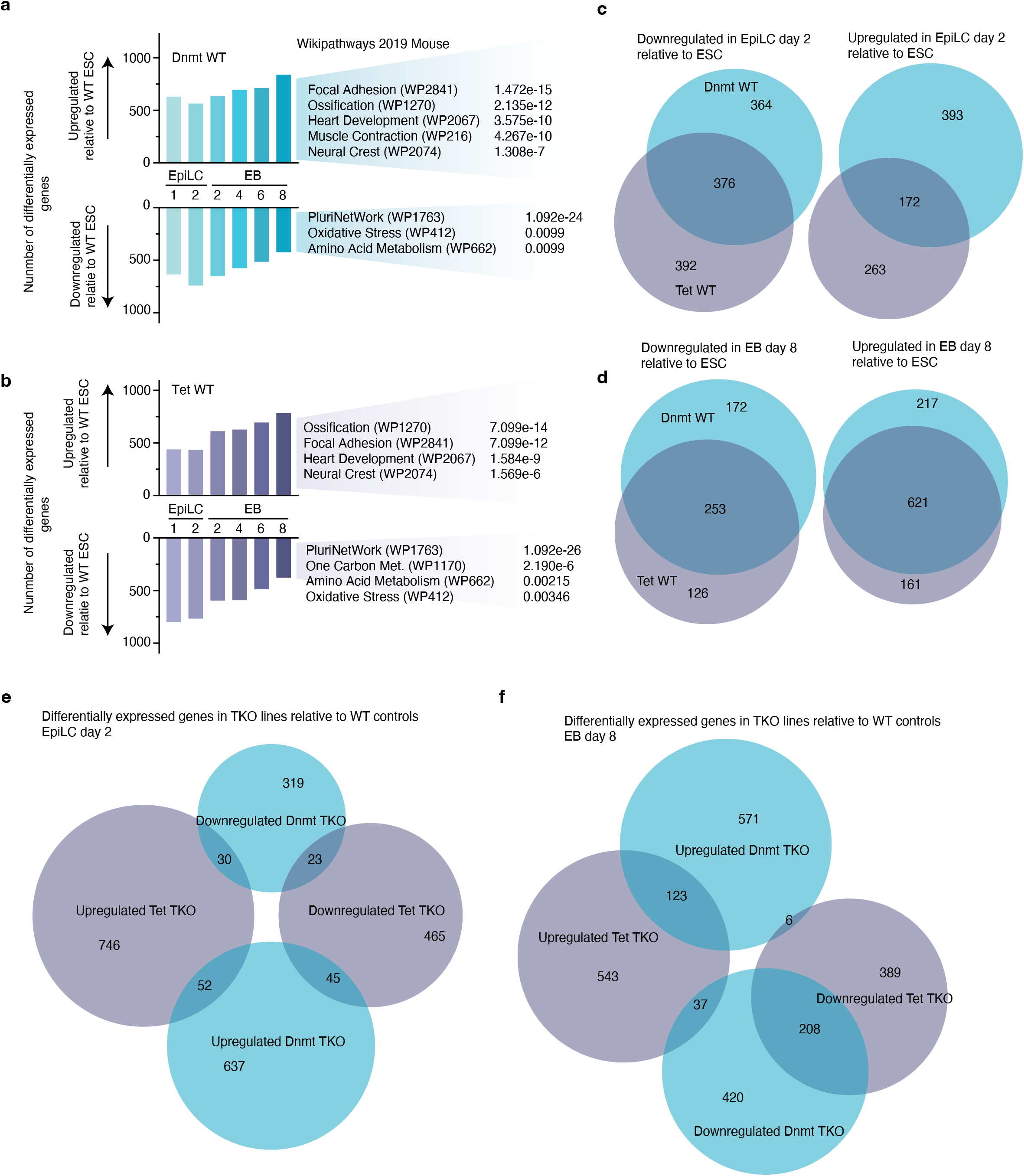
**(a, b)** The number of differentially expressed genes that are significantly upregulated or downregulated (DESeq p<0.05 and dynamic fold change filter) in *Tet* **(a)** and *Dnmt* **(b)** WT EpiLCs and EBs relative to ESCs grown in 2i-LIF (left) as well as gene ontology categories (Wikipathways 2019 Mouse) that differentially expressed genes in day 8 EBs were found to enrich (right) including adjusted p-values. (c) Venn diagrams showing overlap between significantly upregulated and downregulated genes upon EpiLC transition of *Tet* WT or *Dnmt* WT control cells. **(d)** The overlap between significantly upregulated and downregulated genes in day 8 EBs generated from *Tet* WT or *Dnmt* WT control cells. **(e, f)** Venn diagrams showing the numbers of differentially expressed genes in *Tet* and *Dnmt* TKO EpiLCs (e) and embryoid bodies (f) that overlap.

**Supplemental Figure 8 (related to Fig. 6).**
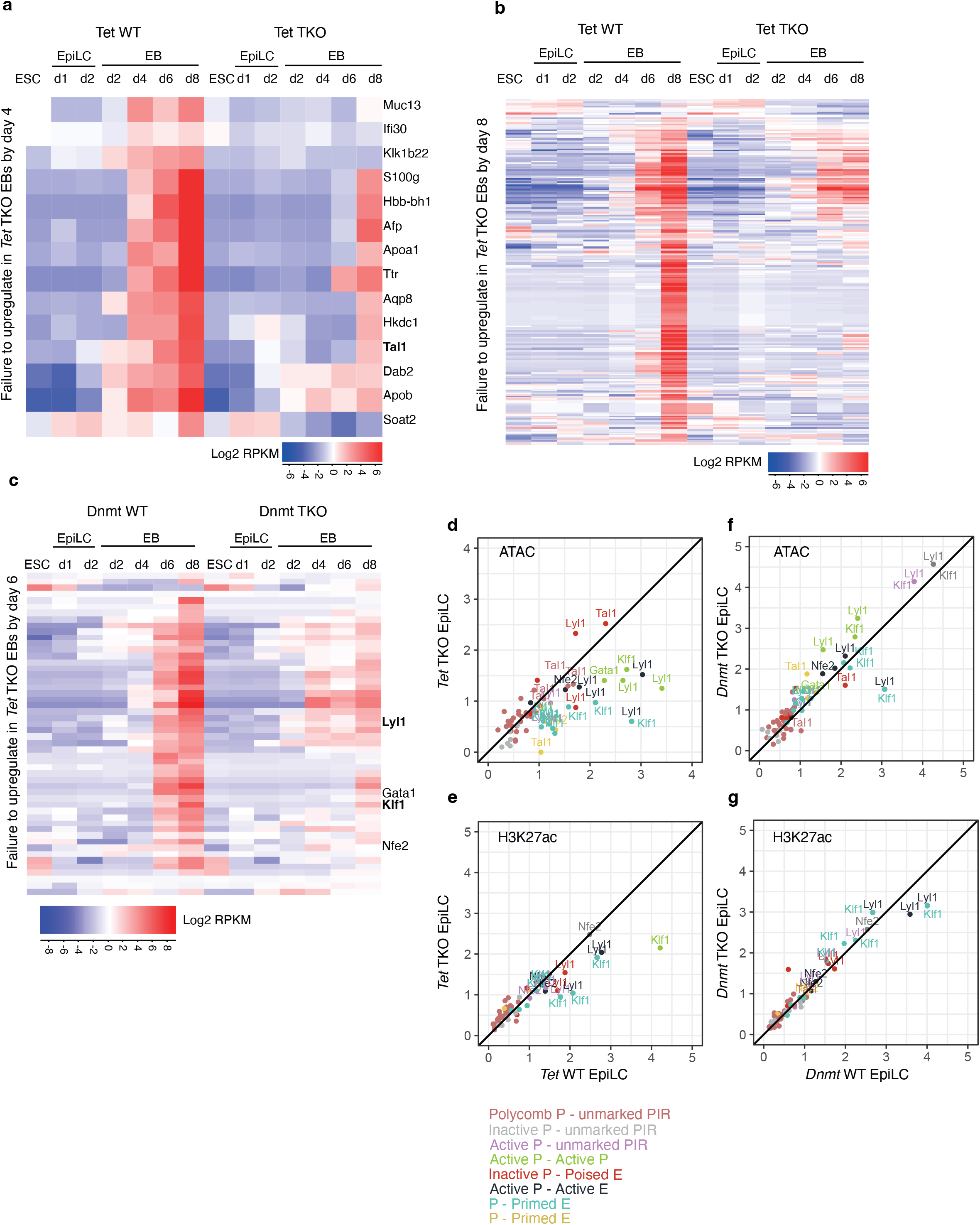
**(a, b)** Heatmap showing genes that are significantly upregulated by day 4 of EB differentiation (a) or day 8 of EB differentiation (b) and remain upregulated in day 8 EBs generated from WT cells, but fail to be upregulated in EBs generated from *Tet* TKO cells (according to DESeq and an dynamic fold change filter). **(c)** Heatmap showing genes that are significantly upregulated by day 6 of EB differentiation (relative to ESCs cultured in 2i-LIF) and remain upregulated in day 8 EBs generated from WT cells, but fail to be upregulated in EBs generated from *Tet* TKO cells (according to DESeq and an dynamic fold change filter). RNA-seq data from *Dnmt* WT and *Dnmt* TKO cells is plotted for these genes. **(d-g)** Scatter plots displaying the levels of accessibility bt ATAC-seq (d, f) or the levels of H3K27ac by ChIP-seq (e, g) in *Tet* TKO (d, e) or *Dnmt* TKO (f, g) EpiLCs relative to respective WT controls at promoter interacting regions (PIRs) involving the indicated blood TFs (labelled in Fig. 6a). ATAC and H3K27ac levels are quantified as reads per million (RPKM). PIRs are labelled by the gene they interact with and coloured by the cluster that the promoter-PIR pair was assigned to (see Fig. 2b).

